# Natural history and ecological effects on the establishment and fate of Florida carpenter ant cadavers infected by the parasitic manipulator *Ophiocordyceps camponoti-floridani*

**DOI:** 10.1101/2022.10.21.513256

**Authors:** Ian Will, Sara Linehan, David G. Jenkins, Charissa de Bekker

## Abstract

1. *Ophiocordyceps* fungi manipulate the behavior of their ant hosts to produce a summit disease phenotype, thereby establishing infected ant cadavers onto vegetation at elevated positions suitable for fungal growth and transmission. Multiple environmental and ecological factors have been proposed to shape the timing, positioning, and outcome of these manipulations.
2. We conducted a long-term field study of *Ophiocordyceps camponoti-floridani* infections of *Camponotus floridanus* ants – the Florida zombie ants. We propose and refine hypotheses on the factors that shape infection outcomes by tracking the occurrence of and fungal growth from hundreds of ant cadavers. We modeled and report these data in relation to weather, light, vegetation, and attack by hyperparasites.
3. We investigated environmental factors that could affect the occurrence and location of newly manipulated ant cadavers. New cadaver occurrence was preferentially biased toward epiphytic *Tillandsia* bromeliads, canopy openness, and summer weather conditions (an interactive effect of temperature, humidity, and precipitation). Furthermore, we suggest that incident light at the individual cadaver level reflects microhabitat choice by manipulated ants or selective pressure on cadaver maintenance for conditions that improve fungal survival.
4. We also asked which environmental conditions affect fungal fitness. Continued fungal development of reproductive structures and putative transmission increased with moist weather conditions (interaction of humidity and precipitation) and canopy openness, while being reduced by hyperparasitic mycoparasite infections. Moreover, under the most open canopy conditions, we found an atypical *Ophiocordyceps* growth morphology that could represent a plastic response to conditions influenced by high light levels.
5. Taken together, we explore general trends and the effects of various ecological conditions on host and parasite disease outcomes in the Florida zombie ant system. These insights from the field can be used to inform experimental laboratory setups that directly test the effects of biotic and abiotic factors on fungus-ant interactions or aim to uncover underlying molecular mechanisms.

## INTRODUCTION

Fungi in the genus *Ophiocordyceps* infect and change ant behavior in a parasite-adaptive manner, with many different host and parasite species interacting across the globe (Araújo et al., 2018; de Bekker, Quevillon, et al., 2014; Evans et al., 2018; Kobmoo et al., 2012; Sakolrak et al., 2018). The common thread among ant-manipulating *Ophiocordyceps* is a terminal summiting behavior whereby the fungus induces the ant to bite and affix itself to vegetation at an elevated position that promotes fungal growth and transmission (Andersen et al., 2009; Evans & Samson, 1984; Hughes et al., 2011). However, specific *Ophiocordyceps*-ant species interactions do vary across climates and habitats, producing the fatal change in host behavior and subsequent fungal development in slightly different ways (Araújo et al., 2018, 2020; Evans et al., 2011; Loreto et al., 2018). How manipulated phenotypes are expressed in different host and environmental contexts could be dependent on the host and parasite species involved, but also be a result of specific environmental pressures experienced at evolutionary and individual timescales.

Variation in “zombie ant” biting substrate, summit height, and developmental timing have been hypothesized to relate to factors such as seasonal changes in habitat and microclimatic light, humidity, or temperature (Andriolli et al., 2019; Cardoso Neto et al., 2019; Chung et al., 2017; Hughes et al., 2011; Lavery et al., 2021; Loreto et al., 2018). Manipulated ant cadavers generally cluster in “graveyards” at higher densities than the surrounding habitat (Andersen et al., 2012; Andriolli et al., 2019; Pontoppidan et al., 2009). Moreover, cadaver distributions and abundances within graveyards can shift over time (Pontoppidan *et al.* 2009), likely reflecting changing environmental factors. Ultimately, *Ophiocordyceps* must transmit to new hosts, requiring the formation of a spore-containing perithecium, or “fruiting body.” Although, the presence of a fruiting body itself does not always indicate active spore dispersal, it is a critical step in spore production and transmission, and, therefore, fungal fitness (Andersen et al., 2012). *Ophiocordyceps* is likely to die, and transmission is halted when the cadaver, or its biting vegetation, detach from their position. Previous studies demonstrated that fruiting bodies are not produced when the cadaver is placed under altered conditions such as other locations in the forest or ant nests (Andersen et al., 2009; Loreto et al., 2014). Moreover, despite performing many infection experiments and collecting multiple naturally manipulated ant cadavers, we have yet to observe fruiting body formation under constrained laboratory conditions (personal observations), which further suggests an environment-fitness link. In addition to abiotic effects, adverse biotic interactions such as hyperparasitism, during which one organism is parasitized by another parasite (Kirk et al., 2008), could hamper *Ophiocordyceps* development and transmission capacity (Andersen et al., 2012; Araújo et al., 2022 in press; Evans, 1982). Fungal hyperparasitisms of *Ophiocordyceps* have regularly been observed by field scientists who study and document the biodiversity of these fungus-ant interactions in the wild (Andersen et al., 2012; Araújo et al., 2020, 2022 in press; Kobmoo et al., 2012). Nevertheless, official species descriptions and reports of these fungi are hard to come by and their potential ecological impact (e.g., impact on trophic cascades) has, to our knowledge, not been documented in any official capacity besides the work that we present here. Moreover, the present study seeks to connect environment to cadaver accumulation and parasite transmission over time. Such a characterization has not yet been reported for the ant host and fungal parasite species commonly associated with Florida, USA. Furthermore, the amount and diversity of data that we collected on natural infections without experimental interference, and within a single study, appears to be uncommon in the reports currently available from other *Ophiocordyceps* species. This, while such natural history-focused observational studies have the power to reveal interesting, biologically relevant patterns that can be tested experimentally.

As such, we tracked the occurrence and development of cadavers from carpenter ants (*Camponotus floridanus*) infected and manipulated by the fungus *Ophiocordyceps camponoti-floridani* across multiple graveyards over ca. 400 days. We documented when, where, and under what conditions summited cadavers appeared. We measured cadaver appearance and fate over time in relation to factors previously linked to *Ophiocordyceps*-killed ant cadavers: weather (temperature, humidity, and precipitation), canopy cover (i.e., general habitat light levels), incident light (i.e., individual cadaver light levels), summiting vegetation and height, ant caste, and the presence of two fungal mycoparasites of *O. camponoti-floridani* (Araújo et al., 2022 in press). We monitored the fate of the developing fungi within the cadavers by recording their survival, growth, and transmission capability. Using these data, we characterized and compared the Florida “zombie ants” to previous studies of cadaver dynamics in related species. These natural history records of *Ophiocordyceps* manipulations lead to new hypotheses about environmental factors that contribute to overall trends and variation in *Ophiocordyceps*-ant interactions in Central Florida and other ecosystems. Behavioral manipulation by *O. camponoti-floridani* and other species is actively studied at the molecular and cellular level to understand the underlying mechanisms (de Bekker et al., 2015; de Bekker, Ohm, et al., 2017; de Bekker, Quevillon, et al., 2014; Fredericksen et al., 2017; Loreto & Hughes, 2019; Mangold et al., 2019; Wichadakul et al., 2015; Will et al., 2020). Biologically relevant interpretations and continuations of these functional investigations will benefit from a better understanding of the accompanying behavior, ecology, and disease dynamics in a natural setting.

## METHODS

### Field sites

We collected data on *O. camponoti-floridani*-infected C. *floridanus* from three wilderness areas in Seminole County, Florida, USA between October 2018 and November 2019: 1) Black Hammock Wilderness (BH), 2) Chuluota Wilderness (CH), and 3) Little Big Econ State Forest (LB). Sampling and data collection were permitted by the Florida Department of Agriculture and Consumer Service’s Florida Forest Service and the Seminole County’s Leisure Services Department, Greenways and Natural Lands Division. We scouted each wilderness area to find *Ophiocordyceps* graveyards with few to no observable ant cadavers in the forest between them. Our aim was not to rigorously compare graveyard versus non-graveyard conditions, but rather to use these high-density areas as an efficient means of finding as many cadavers as possible for our study. Initial cadaver densities during plot establishment at our sites were nearly eight-fold less than previously reported for *Ophiocordyceps* graveyards in Southern Thailand (Pontoppidan et al. 2009), which likely reflects differences in the fungal and ant species studied (*Ophiocordyceps camponoti-leonardi* and *Colobopsis leonardi*), as well as habitat (tropical vs. subtropical). For each wilderness area, we established three graveyard plots of nine-by-nine meters for a total of nine plots. Within a given wilderness area, the identified graveyards were on average 0.39 km apart, having the following GPS coordinates: Black Hammock graveyard 1 (BH-1) N 28°41.993’ W 081°09.522’, BH-2 N 28°42.037’W 081°09.505’, BH-3 N 28°42.459’ W 081°09.484’, CH-1 N 28°37.271’W 081°03.584’, CH-2 N 28°37.211’W 081°03.733’, CH-3 N 28°37.124’W 081°03.545’, LB-1 N 28°40.977’ W 081°09.408’, LB-2 N 28°40.917’ W 081°09.401’, and LB-3 N 28°40.658’ W 081°09.636’ (Fig. S1A in the Supporting Information).

### Ant cadaver data collection

We collected a diverse range of data over time to identify potential drivers of cadaver occurrence and placement, and, subsequent fungal development and transmission. We monitored graveyards over the course of 411 days with a median interval of 60 days between visits to the same graveyard (range 7 to 176 days, except one interval of 232 days). As data collection was labor intensive, different study plots were visited on different days. For each graveyard, we surveyed the entire area from the ground up to ca. 2.6 m above the forest floor for the presence of *Ophiocordyceps*-infected ant cadavers. We thoroughly searched each graveyard for manipulated *C. floridanus* ants during an initial survey to establish a baseline for the number of cadavers present at the beginning of the study. We assigned each cadaver a unique ID, tagged them with a water-resistant label, and recorded their approximate position within the graveyard. We characterized cadavers by their biting vegetation, ant caste, height above the forest floor, and, for a subset, incident light levels on the cadaver. Biting vegetation was classified based upon visual inspection. We determined caste by the cadaver head size; “majors” have a distinct large head and mandibles compared to their “minor” counterparts (Hansen & Klotz, 2005). We measured height (cm) using a tape measure from the cadaver to the forest floor. To estimate incident light (lux) at each cadaver, we used an U12-012 HOBO data logger and software (Hoboware 3.7.21, ONSET), facing up toward the canopy. We averaged three light measurements per cadaver taken over 90 sec held at three different positions directly adjacent to the cadaver. To standardize light intensity, we only took measurements within one hour of solar noon on days with 0% cloud cover. As such, light was not always recorded for each cadaver at first discovery. Because of this necessary delay, we assumed that, being in a subtropical forest, incident light at cadavers was not subject to large changes between manipulated biting and our measurement that would substantially bias the data shadier or brighter than initially experienced by the manipulated ant. We were not able to measure incident light of all cadavers before the COVID-19 pandemic interfered with this project and prevented light measurements at CH-1 and 3 and LB-2 within a reasonable timeframe. In subsequent visits we tagged and characterized newly appearing cadavers and recorded the presence or absence of previously tagged cadavers. We also recorded the presence or absence of a fungal fruiting body. The fruiting body is a necessary step in the *Ophiocordyceps* life cycle for production of sexual spores and transmission. As such, it served as a practical proxy for fungal reproduction capacity (Fig. 1A). Lastly, we recorded cases of hyperparasitism on *Ophiocordyceps* by the fungi *Niveomyces coronatus* (Fig. 1B) and *Torrubiellomyces zombiae* (Fig. 1C) (Araújo et al., 2022 in press). Cadaver photography (Fig. 1) followed the methods of Dal Pos & Rousse, 2018.

**Figure 1:**
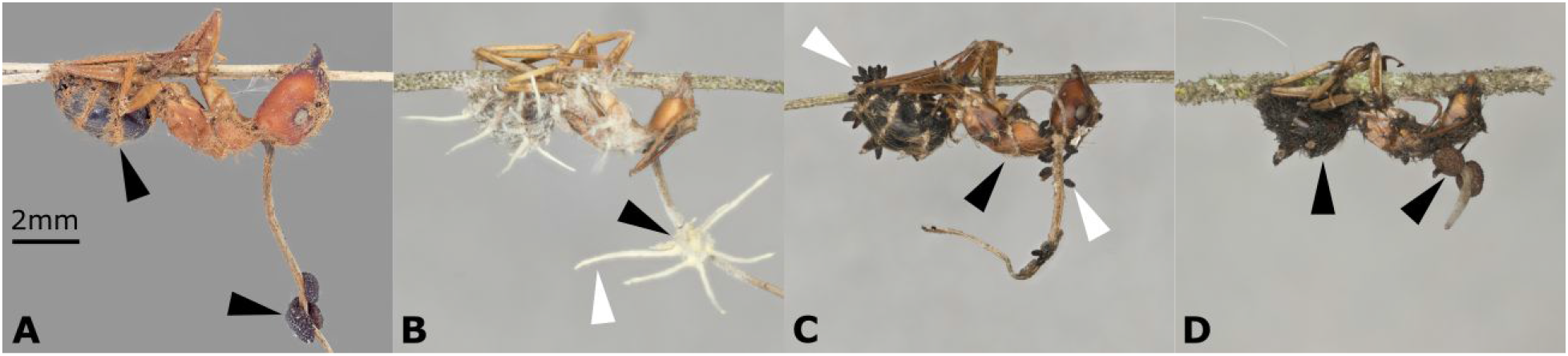
*Ophiocordyceps camponoti-floridani* and mycoparasite morphologies. A) Typical *O. camponoti-floridani* infected cadaver with fruiting bodies (lower black arrow) on the fungal stroma (stalk), mostly found hanging from vegetation in inverted or vertical positions. Hyphae (upper black arrow) often emerge from joints and orifices. B) *Niveomyces coronatus* mycoparasite (white arrow), grown to completely occlude the *Ophiocordyceps* fruiting body (black arrow). C) *Torrubiellomyces zombiae* mycoparasite producing multiple fruiting bodies (white arrows) on top of *Ophiocordyceps* mycelium (black arrow). D) Atypical *O. camponoti-floridani* growth morphology with highly melanized mycelium (left black arrow) covering the cadaver’s body and stunted stalk production, only found in the graveyard with the sparest canopy cover and likely induced by high-light conditions. These cadavers were still able to produce fruiting bodies (right black arrow).

### Graveyard habitat characterization

To describe habitat conditions possibly related to cadaver abundance and distribution within graveyards, we measured vegetation and canopy openness. These surveys were conducted in July 2020 after all cadaver observations had been completed to 1) assure that we did not unnecessarily disturb the plots during our longitudinal study and 2) due to interference from the COVID-19 pandemic. Since the predominant mesic hammock habitat at our subtropical field sites have relatively constant, evergreen canopies (US-FWS, 1999), we considered this single characterization of each graveyard as representative and fair for comparisons among graveyards and sites. To collect vegetation data for each graveyard, we conducted three line-intersect surveys spaced at even intervals perpendicular to the adjacent trail and spanning the 9 m length of the plot. Our line-intersect transects were two-dimensional as we observed variation in vegetation types at different heights. Along the 9 m length of each transect we measured vegetation between the 25^th^ quartile (81 cm) and the 75^th^ quartile value (169 cm) of all cadaver occurrence data. Pieces of vegetation of the same category were measured as continuous if the linear distance in the plane of the transect was ≤5 cm (Caratti, 2006; Klimaszewski-Patterson, 2009). We classified vegetation types to reflect broad categories of plant material found as biting substrates in our cadaver observations and potentially related to functional differences in manipulated ant biting and cadaver retention: *Tillandsia* (epiphytic ball and Spanish “mosses”*Tillandsia recurvata* and *Tillandsia usneoides*, respectively), palmettos (*Arecaceae*), pines (*Pinaceae*), “other woody” plants (64% live oaks, *Fagaceae*) and “other herbaceous” plants (Fig. S1B). Finding many cadavers on *Tillandsia* led us to hypothesize that these plants were an important substrate for manipulated biting. Pine needles are especially slender and prone to dropping off trees, thereby displacing cadavers from their initial perch. Palmettos are notably tougher than many other herbaceous plants in the area and carry distinct slender frays; both the frays and blades of the plant were found as biting substrates, with most cadavers on the palmetto blades. We used “other woody” as a catch-all for all non-pine woody plants that we measured and only observed biting cadavers on twigs of these plants (never on leaves). Finally, the “other herbaceous” category was a catch-all for herbaceous plants other than *Tillandsia* and palmettos; cadavers were not found on leaves but rather slim stems and filaments.

Ball and Spanish mosses are a major substrate for ant biting (see Results section), but they are relatively small and patchy epiphytic plants that could be prone to estimation errors with our line-intersect transects. We verified line-intersect results with targeted *Tillandsia* measurements of random trees within our plots. Trees were selected by dividing each plot into one-by-one meter squares, randomly picking five of the nine squares, and within each square, selecting the centermost tree to measure all present *Tillandsia* up to a height of 2 m. This limit approximated the maximum cadaver height observed in the study (2.6 m) while allowing accurate measurements by our field crew. We measured every *Tillandsia* patch in three dimensions: the maximum height, width, and depth. As in the line-intersect transect data collection, vegetation was counted as continuous when gaps were ≤ 5 cm. Random plot data suggested that the transect method did adequately estimate *Tillandsia* coverage within a graveyard (Fig. S1C). Additionally, *Tillandsia* volume did not simply reflect tree size (no significant correlation with tree circumference at 1 m height and *Tillandsia* coverage; Kendall’s correlation test, p > 0.05).

To quantify canopy cover of each plot, we placed a GoPro Hero 6 on the forest floor facing upward in each of the four corners and in the center of each plot to capture one canopy picture per second, for 20 sec. The GoPro camera was used at wide settings, ISO 100-400 and a shutter speed of 1/500 sec. Of the 20 pictures taken, we selected the sharpest image with the least amount of visible wind disturbance for analysis. We manually adjusted each 4,000 pixel × 3,000 pixel picture to a binary black and white image using Fiji (v 1.53) (Schindelin et al., 2012). Adjusted parameters were within the following ranges: peak hue at 230, saturation between 100-230, and brightness between 87-206. We subsequently set the RGB thresholds to 255 (i.e., white color) to calculate the area of white pixels and determine the amount of open canopy. By averaging the white area from each of the five pictures per graveyard, we calculated the percent open canopy.

### Weather data

We obtained weather data from the National Centers for Environmental Information (NCEI), Land-Based Station WBAN:12815, ca. 32 km from our study sites at N 28°26.000’ W 81°19.000’ (Fig. S1A). This weather station collects daily data on average, maximum and minimum temperature, average relative humidity, and precipitation. We imputed 0.000254 mm for “trace” precipitation records (i.e., 0.00001 inches for the raw data recorded on the imperial system by the weather station) (Akyüz et al., 2013). Such “trace” records indicate that the station detected precipitation, but the amount was below the 0.254 mm measurement threshold.

### *Sanger sequencing of* Ophiocordyceps *ribosomal short subunit*

To confirm that cadavers with atypical fungal morphology at graveyard CH-1 were indeed colonized by *O. camponoti-floridani* (see Results) (Fig. 1D), we verified ribosomal short subunit (SSU) sequence data against the genome of *O. camponoti-floridani* (GenBank WGS accession JAACLJ000000000) (Will et al., 2020). We removed melanized mycelium from a single manipulated *C. floridanus* specimen and used it for DNA extraction following the methods detailed in Will et al., 2020. Purified DNA was subsequently used as a template for PCR using primers NS1 and NS4 (White et al., 1990). We performed PCR reactions with Phusion High Fidelity Polymerase and reagents (New England Biolabs) using a Proflex Base thermocycler (Applied Biosystems) and the following protocol: initial denaturation at 98 °C for 30 sec, 30 cycles of 98 °C for 10 sec, 49 °C for 30 sec, 72 °C for 30 s, and final elongation at 72 °C for 10 min. Sanger sequencing of the amplicon (GenBank accession ON495684) was performed by Eurofins.

### Study overview

A total of 874 *C. floridanus* ant cadavers were surveyed, with 456 of these cadavers newly appearing in graveyards after the study began. Our analyses focused on new cadavers to ensure we had necessary temporal and environmental data to contextualize these observations.

Our study sites were selected in part to capture some of the natural variation in Central Florida habitats, which is reflected in our vegetation transect data (Fig. S1B). We found that only three graveyard-by-vegetation combinations (of the total of 45 combinations) resulted in extreme values using ±1.5 interquartile range (IQR), i.e., possible outlier values typically highlighted by box-and-whisker plots. These graveyard plots were BH-1 and CH-2, with high pine abundances (as many plots lacked pines), and LB-3, with high “other woody” abundance. Each graveyard was also scored for canopy openness using image analysis, thereby generally characterizing the light intensity of the graveyard. The median canopy openness across graveyards was 18.2% and the range was 6.8% to 33.0% with CH-1 being an apparent outlier at 41.3% (> +1.5 IQR).

Each of the nine graveyards showed continued fungal transmission during the study period. The 456 new cadavers were well distributed among graveyards without extreme values as assessed using ±1.5 IQR (range = 6% to 19% of total cadavers from a given graveyard, median = 11%), which corresponded to 29%, 35%, and 36% at the park level, BH, LB, and CH, respectively. Of these new cadavers, 94% belonged to the minor worker caste and 6% to the major worker caste (n = 421 and 29, respectively, six lacked caste records).

Leveraging this longitudinal data set of several hundred ant cadavers, we performed multiple analyses to better understand when and where new *Ophiocordyceps*-infected ants appear and how fungal survival and transmission may be shaped by the environment. Given the relatively well distributed occurrence of new cadavers and variation in vegetation abundances, we combined the data from all graveyard plots across wilderness areas. As such, we performed our analyses on the study-wide dataset, and only singled out graveyard CH-1 for increased scrutiny in relation to an atypical *O. camponoti-floridani* growth phenotype found specifically at that graveyard (see Results).

### Model outlines and rationale

Here, we outline models tested and our rationale in creating them. The complete reporting and final model selection can be found in the relevant Results sections below.

#### Model-1

To determine if vegetation type and/or the abundance of that vegetation influences the relative accumulation of newly manipulated ant cadavers, we used a binomial generalized linear mixed model (GLMM). We generated model hypotheses of the proportion of cadaver counts within a plot by interactive or additive fixed effects of vegetation type and relative abundance of that vegetation within the plot. Cadaver counts were combined into study-long totals, per graveyard plot, and per vegetation type. To account for spatial pseudo-replication, we tested the introduction of random effects for graveyard plots and wilderness areas in the models. In cases where we could not converge a model with any random effect, we introduced graveyard plot as a fixed effect. The full model described the number of cadavers (dependent variable) by the interactive fixed effect of vegetation type and abundance, with a random effect of graveyard plot nested within wilderness area (independent variables).

#### Model-2

To investigate if cadaver height indicated an adjustment to or selection by light levels, we constructed GLMMs using the Gamma distribution for our slightly positively skewed height data. We tested height as a response to canopy openness, incident light, or both variables. Given the lower sample size and gaps in coverage, most models could not be fit as GLMMs with random effects of wilderness area or graveyard plot. In such cases, we tested generalized linear models (GLMs) using plot or wilderness area as a fixed effect. What we considered to be the full model described cadaver height (dependent variable) by additive fixed effects of canopy openness, incident light at the cadaver, and wilderness area (independent variables).

#### Model-3

To model the accumulation of new cadavers in plots over time in relation to habitat and weather variables, we used Poisson GLMMs. Each visit to a graveyard was scored for the number of new cadavers since the last visit and averaged weather conditions during that interval. Weather station records included temperature, humidity, and precipitation. We tested the mean daily humidity and mean daily precipitation of a visit interval in relation to the number of cadavers found. For temperature, we could have constructed this interval mean value using daily maximum, minimum, or mean temperatures. We had no *a priori* reason to expect any one temperature measurement (or the difference between daily minimum and maximum) to be the most related to cadaver occurrence. We selected daily maximum temperature based on preliminary linear models and plotting (data not shown), although no one temperature measurement presented a much clearer relationship to cadaver rate than any other.

In addition to weather variables, we used our habitat characterization data to determine the effect of vegetation on new cadaver accumulation. Because *Tillandsia* were favored for new cadaver occurrence (see Results, *Model-1*), we included *Tillandsia* coverage of graveyards, using two possible metrics. First, we described *Tillandsia* coverage as a percent coverage relative to other vegetation types within a plot, using transect line-intersect data. Second, we used our random tree measurements to test absolute *Tillandsia* abundance. We also included canopy openness in our models because it was our only ubiquitously collected light metric and a potential proxy for various graveyard conditions dependent on canopy cover. Although, in principle, graveyards were nested within wilderness areas, models incorporating a nested random effect failed to converge with our data. As such, we used graveyard only. To account for time between visits, we included an offset term for days since last visit, as the number of new cadavers found in a graveyard would intuitively scale with time.

Taken together, this resulted in full models of new cadaver accumulation (dependent variable) that included the interactive effect of weather data (precipitation, humidity, and daily maximum temperature), a *Tillandsia* coverage measurement (either abundance or relative abundance to other vegetation), canopy openness, and a random effect of graveyard plot with an offset for visit-interval length (independent variables).

#### Model-4

We employed binomial GLMMs to investigate fungal transmission capability (i.e., fungal fruiting body formation). We tested weather variables as we did for cadaver accumulation (see above, *Model-3*). Each cadaver was related to the averaged weather data from the interval between when it was last seen (or the plot was last visited) and observation of fruiting or detachment. We also tested cadaver height, biting vegetation type, and graveyard plot canopy openness. We accounted for repeated sampling in space with a random effect of graveyard plot and for repeated temporal effects with a random effect of time, represented as date-bins for our visits to the field. Bins were based on the time between the first and last cadavers observed to reach a transmission outcome (either fruiting or detachment) divided into the maximum number of bins that contained at least three cadaver observations per bin. This produced 11 visit bins of 29 days each. All but two visit bins had observations from multiple plots. As such, this final model described fungal fruiting or detachment (dependent variable) by a three-way weather interaction (temperature, humidity, and precipitation), cadaver height, vegetation type, and canopy openness fixed effects, and random effects of both graveyard plot and time (date-bins) (independent variables).

### Data analysis and statistics

The environmental variables collected in this study represent informed explorations, based on previous *Ophiocordyceps* research, that potentially shape the life history of *Ophiocordyceps*-killed ant cadavers (Andriolli et al., 2019; Cardoso Neto et al., 2019; Chung et al., 2017; Hughes et al., 2011; Lavery et al., 2021; Loreto et al., 2018). When our objective was to only report natural history observations, we did not statistically analyze data. However, where appropriate to test hypotheses, we performed statistical tests using R (v 3.6 and 4.1) through R Studio (v 2021.09.2) (R Core Team, 2021; RStudio Team, 2015). Generalized linear mixed models and GLMs were fit using R package lme4 (v 1.1-27.1) or glmmTMB (v 1.1.2.3) (Bates et al., 2015; Brooks et al., 2017). When fitting models, we scaled all continuous variables to standardized variables with different ranges and units. Competing model hypotheses were compared with the corrected Akaike information criterion (AIC) using R package performance (0.8.0) to arrive at our final model and significant effects (Akaike, 1974; Lüdecke et al., 2021). Model selection using AIC has several philosophical and practical advantages because it 1) is consistent with multiple working hypotheses (Elliott & Brook, 2007), 2) balances goodness of fit with model complexity to identify the most efficient model among those being compared (Burnham & Anderson, 2002), and 3) is asymptotically equivalent to leave-one-cluster-out cross validation (Fang, 2011). Model comparisons emphasized AIC weight (*w_i_*; the probability that a model is most efficient among those listed) and δAIC, where values > 2 indicate clear model choice (Burnham & Anderson, 2002). We report AIC and effect estimates for tested models with better fits than a null model. The most plausible models (by AIC) were evaluated for adherence to model assumptions and fit by pseudo-R^2^_GLMM_ and Nagelkerke’s R^2^ scores using R package performance (Lüdecke et al., 2021; Nagelkerke, 1991; Nakagawa & Schielzeth, 2013). All plots were generated using R package ggplot2 (v 3.3.5) (Wickham, 2016).

## RESULTS

### Cadavers are most often found on Tillandsia (Model-1)

Cadavers were found most often on *Tillandsia;* 71% had attached themselves to *Tillandsia*, 14% to twigs of other “woody plants”, 10% to pine needles, 3% to “other herbaceous” plants, and 1% to palmettos (n = 318, 64, 45, 13, and 5 cadavers, respectively, 11 lacked vegetation records). As most of the ant cadavers across the graveyards were attached to *Tillandsia*, we asked if vegetation type was predictive of cadaver distribution within a plot. Alternatively, cadaver attachment could be random and simply correlated to vegetation type abundance. The model we used to investigate this relationship described the relative abundance of cadavers by the type and/or abundance of their biting vegetation (Model-1, see Methods). Model selection by AIC favored the full model with interactive fixed vegetation effects, and with a nested random effect of graveyards within wilderness areas (Tables S1-S2). The marginal pseudo-R^2^_GLMM_ for fixed effects was 0.55, and the conditional pseudo-R^2^_GLMM_ including random effects was 0.64. This final model identified the strongest effect on cadaver distribution as an interaction term indicating the relative abundance of epiphytic *Tillandsia* (β = 30.45, p < 0.001) (Fig. 2). This effect is relative to “other woody” as the intercept, which was typically the most abundant vegetation in our study sites (Table S2). We also found the abundance of pines (β = 5.47, p < 0.001) to have positive effects on cadaver accumulation while the “other herbaceous” vegetation category had a modest negative effect (β = −1.53, p = 0.032).

**Figure 2:**
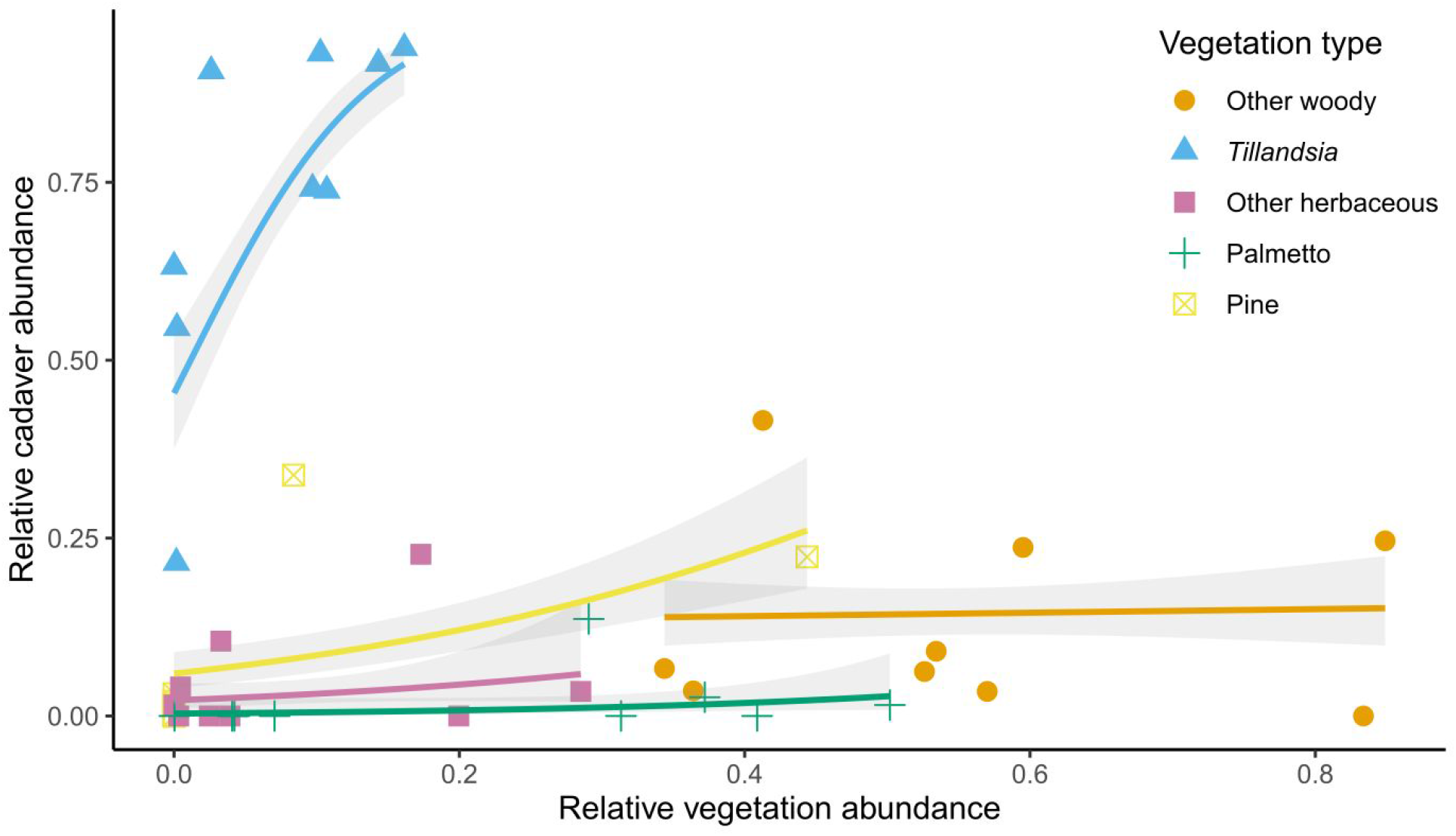
Distribution of cadavers across vegetation types strongly favor *Tillandsia* in a density dependent manner. A GLMM indicated the strongest positive predictor of where cadavers appeared within a graveyard to be the relative abundance of *Tillandsia* (β = 30.45, p < 0.001). Additionally, the abundance of pines positively predicted cadaver abundance (β = 5.47, p < 0.001). Cadavers were less likely to appear on the “other herbaceous” type vegetation (β = −1.53, p = 0.032). Vegetation type is shown as follows: “other woody” (orange circles), *Tillandsia* (blue triangles), “other herbaceous” plants (purple squares), palmettos (green crosses), and pines (yellow crossed squares). The solid lines represent a binomial fixed effects-only GLM per vegetation type in matching color. The gray shaded regions are 95% confidence intervals.

### Cadaver heights correlate to canopy openness but not incident light levels (Model-2)

We measured the height of *Ophiocordyceps*-infected cadavers, finding a median height above the forest floor of 124 cm (range = 12 to 262 cm, n = 446). We hypothesized that summitting height would reflect selection for incident light at the cadaver since evidence from infected *Camponotus atriceps* in Brazil suggests that light affects their height, spatial distribution, and fruiting body formation (Andriolli et al., 2019). As such, we predicted that cadaver height would correlate with canopy openness to maintain cadaver light levels within a preferred range in the shrub and subcanopy layers. Following this reasoning, cadaver heights would decrease under brighter canopies while they would not necessarily correlate directly with incident light levels.

We were only able to collect individual cadaver light readings for a subset of the cadavers and graveyards (n = 115 cadavers, across LB-1 and 3, BH-1, 2, and 3, and CH-2). The median cadaver incident light level was 4044 lux, and the range was 706 to 31,582 with eight additional readings at our light monitor’s maximum level (32,280 lux) (Fig. 3A). Notably, half of the samples occupied a much narrower range (2,152 to 12,332 lux, defined by the 25^th^ and 75^th^ percentiles) creating the prominent peak visible in Figure 3A. For subsequent analysis we only used cadavers with both recorded height and below-maximum incident light (n = 107). The height data for this subset of cadavers was similar to that of the full dataset (range = 21 to 239 cm, median = 110 cm).

**Figure 3:**
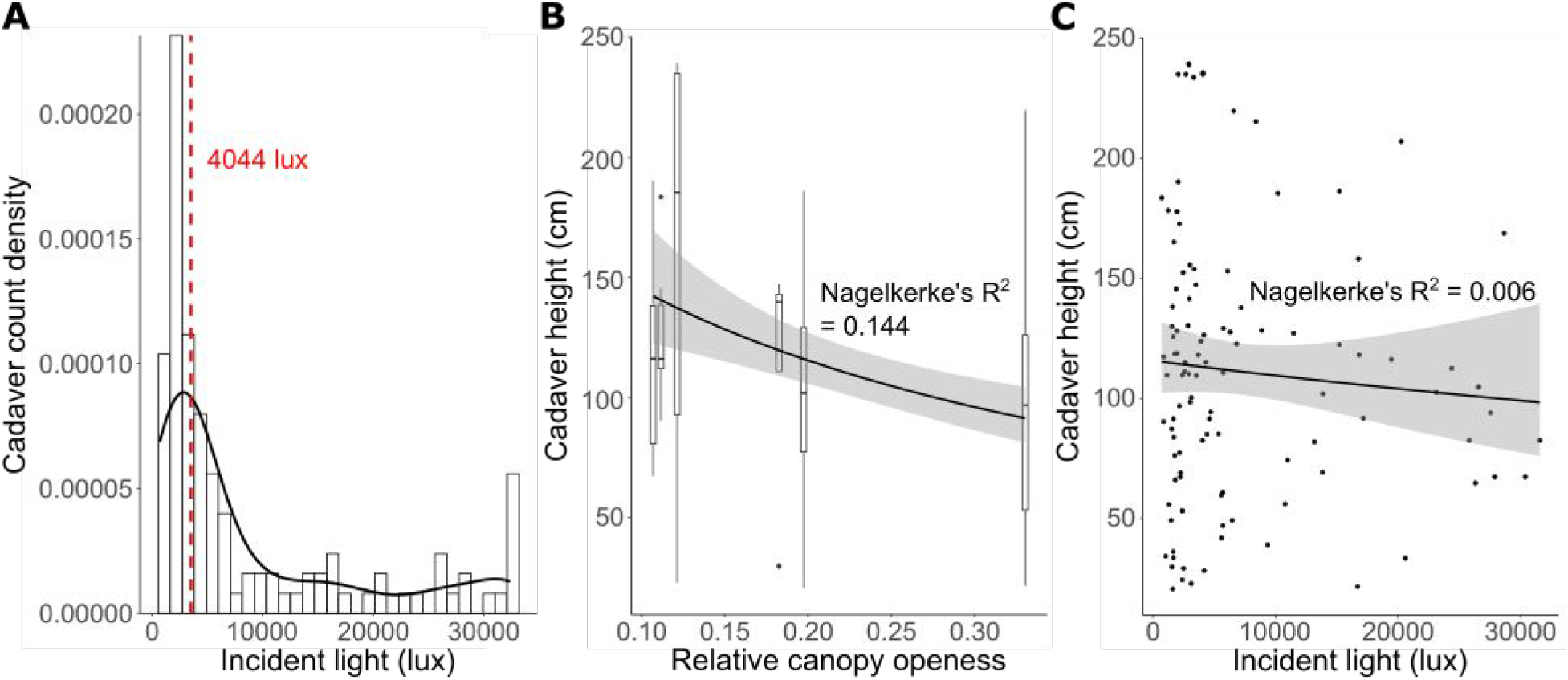
Cadaver height may be a response to canopy openness to maintain incident light levels. A) Distribution of all cadaver incident light data collected (n = 115) shown by a histogram (bars) and density plot (black line). The median value (red dashed line) was 4404 lux. The data contain eight readings at our instrument’s maximum reading capability (32,280 lux). This distribution shows a peak near the median, with 50% of the data between 2,152 and 12,332 lux. The histogram bar conspicuously higher than the rest resides within this peak range, containing 20% of the data within a span that includes 3% of the total range of light measurements (1,759 to 2,811 lux). B) Modeling cadaver height by canopy openness, we found a negative relationship (coefficient = −0.37, p = 0.018). C) Cadaver heights were not related to incident light (coefficient = −0.02, p = 0.679). This suggests that higher overall light levels within a graveyard select for lower cadaver heights, thereby maintaining incident light levels closer to a possible optimum near 4404 lux. Box plots in B) present cadaver height distributions per graveyard, within which all cadavers share the same canopy openness value. Black dots in C) represent individual cadaver measurements. B, C) Black lines are Gamma GLM fits with 95% confidence intervals in the shaded region.

To investigate our hypothesized adjustment (or selection) of cadaver height in response to light levels, we tested models of cadaver height in relation to canopy openness and incident light (Model-2, see Methods) (Tables S3-S4). A canopy openness model (with a fixed effect of wilderness area) was selected by AIC as the most efficient model (Nagelkerke’s R^2^ = 0.144) (Fig. 3B). Canopy openness had a negative effect on cadaver height (coefficient = −0.37, p = 0.018), i.e., cadavers in more open plots were found at lower heights. Models using incident light alone were less efficient than the null model, nor was this effect informative in a model with canopy openness (Tables S3-S4). In line with this finding that canopy openness and incident light had different relationships to cadaver height, we additionally verified that canopy openness and incident light were not strongly correlated to each other (Fig. S2).

### A non-canonical *Ophiocordyceps* morphology found under high canopy openness

*Ophiocordyceps* growth from ant cadavers is typically characterized by limited hyphal growth from between the ant sclerites and joints and a long stalk that carries the fruiting body emerging from behind the head (Fig. 1A). However, at graveyard CH-1, we observed an atypical *Ophiocordyceps* growth morphology (n = 34 of 50 total CH-1 cadavers). The atypical fungal phenotype was characterized by seemingly melanized thread-like mycelium that loosely wrapped the cadaver (Fig. 1D). By extracting DNA from this mycelium and sequencing the SSU region we confirmed that this growth was produced by *O. camponoti-floridani*. The stalks on these cadavers were also much shorter and wider with a dark brown coloration at the tip. Even with the atypical growth form, cadavers still produced fruiting bodies (n = 8), albeit much closer to the cadaver’s head than usual (Fig. 1D). Canopy openness at CH-1 was 41.3%, which was the only extreme canopy openness value (see Methods). Heights were only slightly different between cadavers with atypical morphology, typical cadavers at CH-1, and all other cadavers in the study (median height = 104, 155, and 124 cm respectively). Unfortunately, we lack incident light measurements at CH-1 to further test a relationship between fungal morphology and light intensity.

### Cadaver accumulation rate correlates with weather and season (Model-3)

We investigated if weather helps predict when and where new cadavers appear. We calculated rates of new cadaver accumulation for each interval of time between visits to a graveyard (i.e., the number of new cadavers found divided by the days since the plot was last visited). To generate a study-wide mean rate, we averaged the rate of all plots for every day of the study for which at least half (≥ 5) graveyards had recorded data. Cadaver accumulation rates varied through the study from ca. one cadaver every two weeks (0.07 cadavers/day, 3-8 November 2018) to nearly three-fold higher with one cadaver every five days (0.20 cadavers/day, 18-30 April 2019), with a median one cadaver every seven days (0.15 cadavers/day) (Fig. 4). The rate of cadaver accumulation appeared to peak with seasonal trends in temperature and, possibly, humidity and precipitation (Fig. 4A) warranting further investigation.

**Figure 4:**
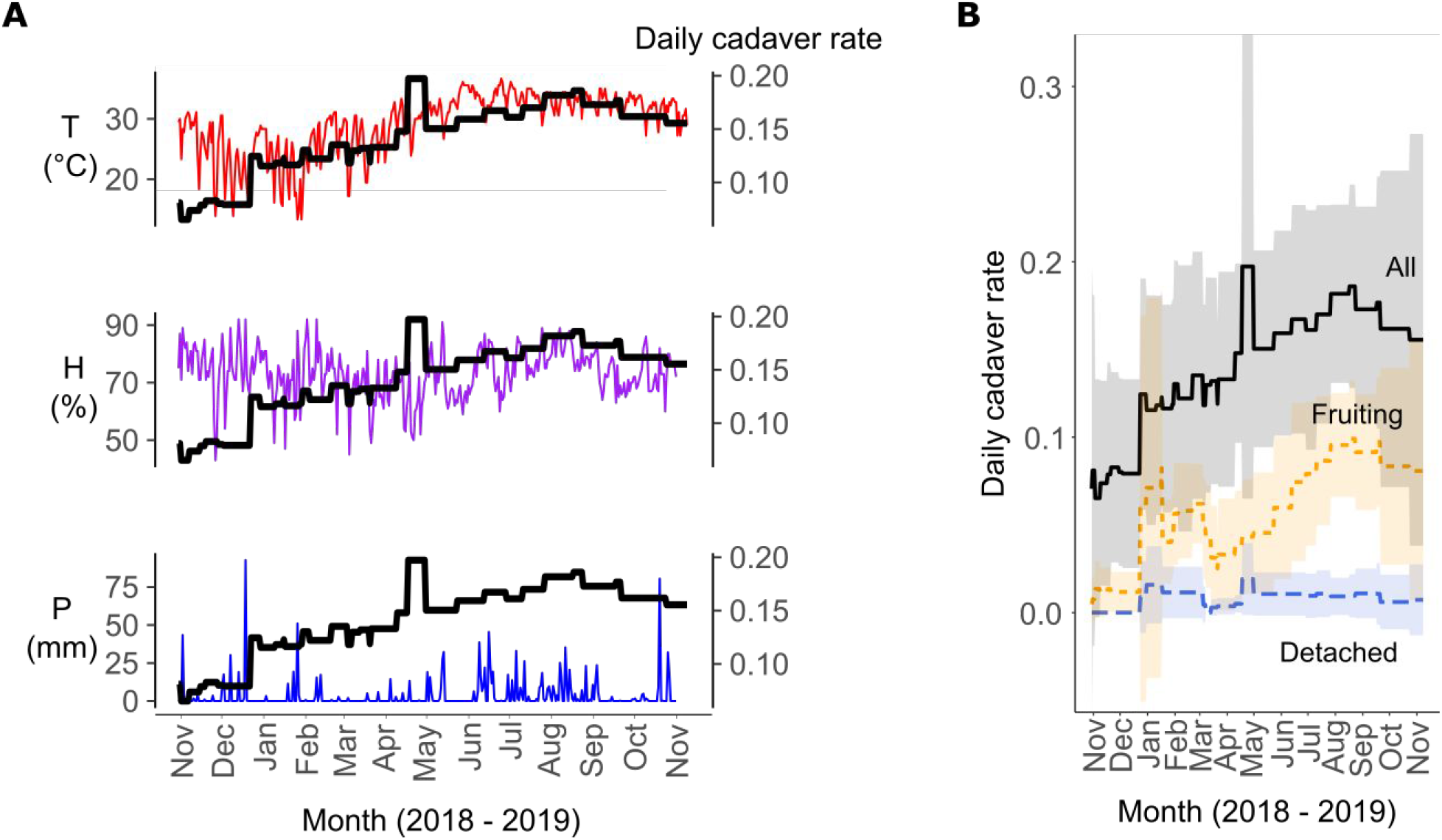
Cadaver appearance and fruiting body formation rates appeared to have warm-weather seasonal trends. A) Estimated daily cadaver rate (black) in relation to weather variables showed a possible trend for a higher rate of cadaver accumulation in the warmer and wetter summer. Cadaver rates are mean values for all study dates with at least five of the nine graveyards having been recorded. A GLMM also highlighted the effects of daily maximum temperature (T, red) (β = 0.28, p < 0.001), daily total precipitation (P, blue) (β = −0.27, p = 0.004), and the three-way interaction of temperature, precipitation, and daily average humidity (H, purple) (β = 0.20, p =0.003). B) Rates of new cadaver appearance (black solid line), cadaver fruiting body formation (orange short-dashed line), and detachment (blue long-dashed line). Rates were calculated for every visit interval per plot as the number of cadavers appearing, forming a fruiting body, or detaching divided by the interval of days between consecutive visits. During the timescale of this study, many more cadavers generated fruiting bodies than detached and this pattern may follow a seasonal trend peaking in the late summer. Shaded areas are 95% confidence intervals.

Model-3 (see Methods) combined weather and habitat data to model new cadaver accumulation rates. As an abbreviated method similar to stepwise selection, we tested a fully interactive effect of weather terms as well as reducing the number of weather terms to only those that were significant in a full model (p ≤ 0.05) and without interactions. We used this approach to select a set of possibly more efficient weather variables than the full three-way interaction of precipitation, humidity, and temperature (Tables S5-S6). We incorporated habitat into the models by combinations of *Tillandsia* abundance and canopy openness. We selected a final model containing significant weather terms initially identified from the full model and canopy openness by AIC comparison. This final model included a three-way interaction of weather terms (β = 0.20, p = 0.003), canopy openness (β = 0.26, p = 0.007), and the random effects of plot and visit-length intervals (marginal pseudo-R^2^_GLMM_ = 0.56, conditional pseudo-R^2^_GLMM_ = 0.75; Table S5). Similar models exchanging the canopy term to describe plot with either a line-intersect or random-tree *Tillandsia* coverage measurement or no such plot habitat term at all were nearly as efficient (Tables S5-S6).

### *Ophiocordyceps* development and transmission in relation to environmental conditions (Model-4)

Given that environmental variables appeared to influence manipulated ant attachment and/or where cadavers are retained in the short-term, we hypothesized that these summiting conditions could be adaptive for fungal parasite fitness outcomes (fruiting or detaching before fruiting), either directly or in advance of key conditions for transmission. Light may be important (Andriolli et al., 2019), as might other environmental conditions. Here we evaluated vegetation, cadaver height, weather data, canopy openness, and time until fruiting body development or cadaver detachment to investigate how environmental conditions could relate to *Ophiocordyceps* transmission outcomes.

Of 445 new cadavers with vegetation data collected, 178 produced a fruiting body within the study interval, of which 8 detached from their biting substrate after producing fruiting bodies. These eight cadavers were analyzed with the attached cadavers because they presumably had an opportunity to transmit spores and infect the next generation of hosts (positive fitness). On the other hand, 26 cadavers detached before a fruiting body appeared and were deemed to not have been able to transmit (no fitness). Although cadavers may detach due to multiple reasons, our model testing sought to determine if environmental conditions are among the factors that could delay fruiting to a degree that the fungus is at risk to detach before transmitting and/or directly promote detachment of the cadaver. Cadavers that remained in the study without reaching either of these transmission outcomes were not included in the following analyses, as their reproductive fates were undetermined.

Given that our sampling schedule varied greatly due to labor-intensive data collection, we could not track times needed for individual new cadavers to produce a fruiting body or detach. However, by using the time interval between observing one of these events and the last sampling visit to that plot, we determined the most frequently spanned number of days between new cadaver occurrence and fruiting body development or detachment (i.e., the mode number of days across this interval for each cadaver). Development of a fruiting body often occurred sometime between 25 and 45 days after a cadaver established (89% of fruiting cadaver observations span this interval) with detachment most often occurring at 64 days (73% of detached cadaver observations include this number of days) (Fig. S3). As most cadavers tracked to a transmission outcome developed fruiting bodies and did so quicker than cadavers that detached, we infer that most cadavers in our graveyards successfully produce fruiting bodies. Although, our initial field observations included some clearly old cadavers (e.g., desiccated, without visibly viable fungal growth) that did not have fruiting bodies but appeared largely intact, suggesting that not every established cadaver will transmit.

We modeled transmission outcomes (i.e., fungal fruiting or cadaver detachment, Model-4, see Methods) with weather data (as above for Model-3), cadaver height and vegetation type, and canopy openness. Our most plausible model (Tables S7-S8) predicted cadaver fruiting or detachment as a function of canopy openness (β = 1.67, p = 0.001) and an interaction between humidity and precipitation (β = 1.44, p = 0.082), with a fairly clear signal (marginal pseudo-R^2^_GLMM_ = 0.45 and conditional pseudo-R^2^_GLMM_ = 0.58; Fig 5). This model included random effects to account for space (graveyard plot) and timing (field visit date-bin, see Methods).

**Figure 5:**
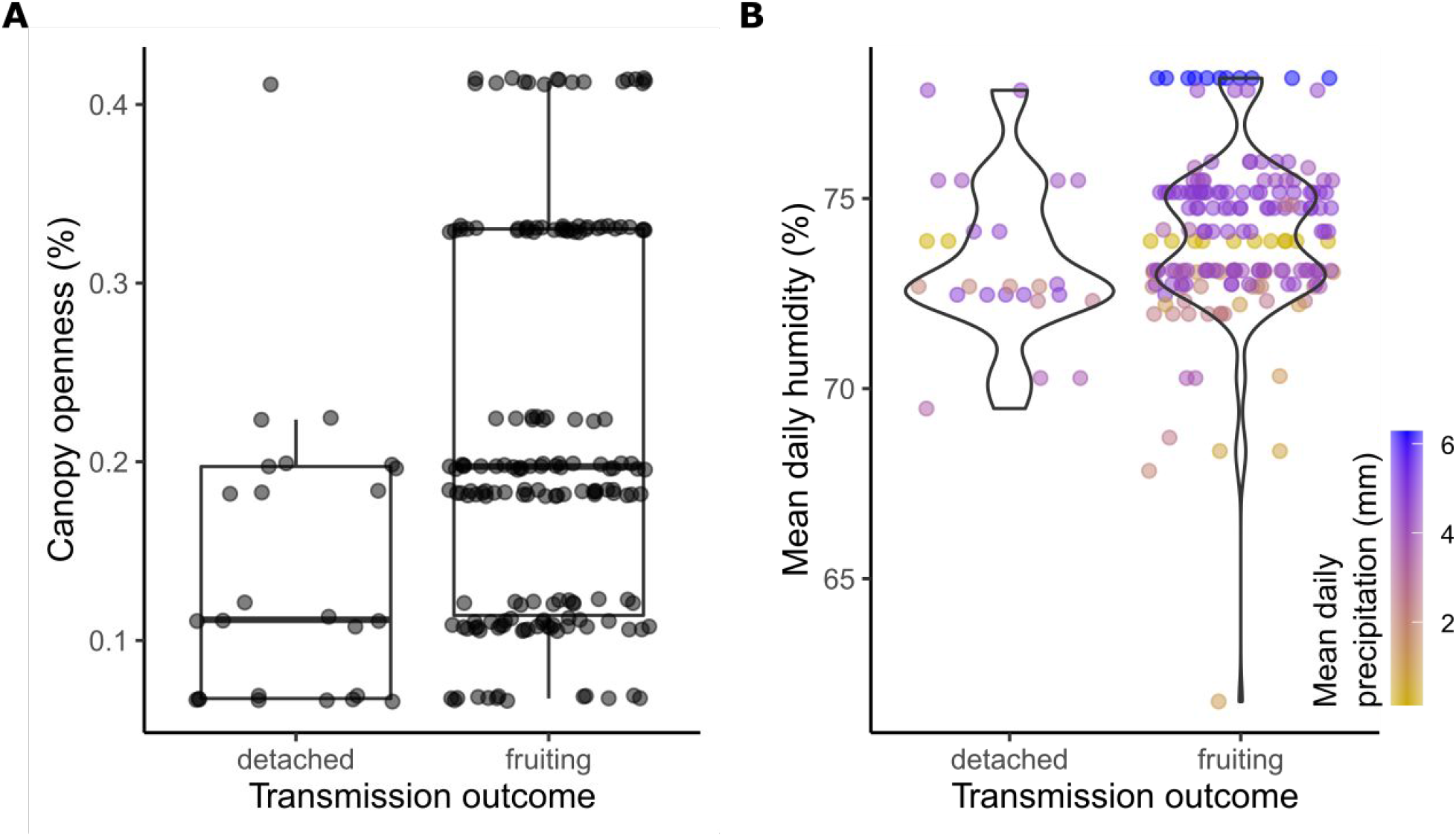
*Ophiocordyceps* transmission may improve in brighter, more open habitat and with wetter conditions. Each dot represents a single cadaver and is A) plotted according to the canopy openness of the plot it was found in or B) the mean daily weather measurements for the interval of time between its last observation and reaching a final transmission outcome (detachment or fruiting). Our most efficient model of cadaver transmission outcome contained the fixed effects of A) canopy openness (β = 1.67, p = 0.001) and B) an interaction of humidity and precipitation (β = 1.44, p = 0.082). Models containing additive effects of humidity or precipitation were less efficient, suggesting that *Ophiocordyceps* produces fruiting bodies most reliably during wet conditions with both high humidity and precipitation.

### Fungal mycoparasites colonize *O. camponoti-floridani*

*Ophiocordyceps camponoti-floridani* is itself parasitized by fungi (*T. zombiae* and *N. coronatus*) (Araújo et al., 2022 in press), which plausibly could affect *Ophiocordyceps* transmission. Of the 456 new cadavers found, 20 (4%) were infected by mycoparasites, 6 by *N. coronatus* and 14 by *T. zombiae* (Fig. 1B and C, respectively). Both mycoparasites were present in each of the three wilderness areas and in some cases co-occurred in the same plot. We did not observe any cadavers co-infected by both mycoparasites.

Based on our sampling of new cadavers, *T. zombiae* colonized *O. camponoti-floridani* most often within 49 days or less (79% of *T. zombiae* infected cadavers span this interval) (Fig. S4A). In three cadavers, *T. zombiae* was only first observed months after they initially attached, suggesting that *T. zombiae* colonizes *O. camponoti-floridani* sometime after ant manipulation rather than competing during the initial phases of infection (Fig. S4A). All six *N. coronatus* infections occurred within 110 days or less (Fig. S4A). However, based on some cadavers having *O. camponoti-floridani* stroma already developed before the onset of hyperparasitism (n = 4), we again surmise that *N. coronatus* attacks established *O. camponoti-floridani*.

While we found *T. zombiae* nearly year-round, the few *N. coronatus* infections were only observed in November (Fig. S4B). However, low sample size and long intervals between observations of new *N. coronatus* infections (110 to 232 days) preclude more definitive statements about possible speed of infection or seasonal trends.

Since both mycoparasite species grow on *Ophiocordyceps* mycelium directly (Araújo et al., 2022 in press) (Fig. 1B and C), we hypothesized that their attack could reduce *O. camponoti-floridani* fruiting body production or function. Of the hyperparasitized cadavers, two produced fruiting bodies after being colonized, six had both a fruiting body and a mycoparasite discovered on the same graveyard-observation day, and the remaining 12 never produced a fruiting body. These 12 cadavers were observed over months (range = 72 to 263 days, median 199 days), far greater than the 25 to 45 days estimated for time until fruiting (see above, Fig. S4A). From this, we infer that *Ophiocordyceps* in these cadavers had died and/or would never transmit. Whether counting only the two cadavers fruiting after hyperparasite infection (10% fruiting rate) or the six cases with unknown order of hyperparasitism and fruiting body formation (40% fruiting rate), hyperparasitism appeared to harshly limit *Ophiocordyceps* transmission in comparison to non-parasitized *Ophiocordyceps* (76% fruiting, n = 178). For this non-parasitized comparison, we accounted for observation time by using non-fruiting cadavers monitored for a similar time as their hyperparasitized counterparts (n = 57, range = 72 to 224 days, median = 110 days).

## DISCUSSION

To better understand ecological conditions of *C. floridanus* ants manipulated by the fungus *O. camponoti-floridani*, we monitored over 400 ant cadavers for more than a year at three subtropical Florida wilderness areas. Environmental conditions or season affect the timing, position, longevity, and development of *Ophiocordyceps* infections (Andriolli et al., 2019; Loreto et al., 2014, 2018). While new cadavers appeared throughout the year, they accumulated more quickly during warmer and wetter months. It is not yet clear whether seasonal weather drives manipulated ant occurrence or if conditions are proxies for other seasonal effects. For example, *Camponotus* colony activity varies with season and resources, which also affects annual timing of mating flights (Hansen & Klotz, 2005). Most simply, the number of new cadavers may correlate to active *C. floridanus* population size as foraging ants become more frequently exposed to *O. camponoti-floridani* spores. However, previous research demonstrates that cadaver distributions are also directly affected by abiotic factors and in some cases possibly more so than by biotic ones, such as proximity to hosts (Loreto et al., 2014; Pontoppidan et al., 2009). Notably, increased humidity acutely promoted spore release in a related *Ophiocordyceps* species (Andersen et al., 2012). While we cannot affirm specific microclimate effects per cadaver based on regional weather pattern data, the moderate correlation between moisture and fruiting body formation that we observed appears to be in line with the link between humidity and infectious spore release that was previously established (Andersen et al., 2012).

Regardless of cadaver timing, strong selection seems apparent for where manipulated ants attach themselves. Although epiphytic *Tillandsia* were relatively rare in our study plots, they held the vast majority of cadavers, beyond expectations based on *Tillandsia* abundance alone.

The slim, bunched stems of the *Tillandsia* (and perhaps pines) may be the easiest attachment sites for manipulated ants. However, *C. floridanus* and other ant species can latch onto tougher and wider substrates such as woody twigs or palmetto blades as evidenced by ant cadavers at our study sites and others (Loreto et al., 2018). Infected ants often display convulsions and uncoordinated movement (de Bekker et al., 2015; Hughes et al., 2011; Will et al., 2020). Speculatively, this may mean *Tillandsia* tangles catch stumbling ants and offer easy access to many suitable locations for final attachment by uncoordinated, infected ants. Another adaptive explanation for preferred attachment or retention to *Tillandsia* could be that its dense tangle of highly flexible stems provides protection from scavengers. In contrast to results on *Tillandsia*, “other herbaceous” vegetation was negatively associated with cadaver abundance. Other herbaceous vegetation seemed at least as pliable and bitable as *Tillandsia* or pine needles. Taken together, this suggests that manipulated ants can attach to a wide variety of vegetation, but actively select or are retained on *Tillandsia* more so than other vegetation.

Beyond summiting vegetation type, light also played an important role in cadaver position. By comparing cadaver heights, canopy openness, and incident light at the cadaver, we found cadavers attached at lower heights under greater canopy openness, but height was not related to incident light at the cadavers. We inferred that manipulated ants climbed less high under more open canopies to reach similar light levels. Furthermore, we found cadavers in this subtropical habitat at heights often greater than those reported for *Ophiocordyceps* species in the tropics (mean height range = 17 cm to 109 cm) (Andersen et al., 2009; Andriolli et al., 2019; Cardoso Neto et al., 2019; Hughes et al., 2011; Lavery et al., 2021). We hypothesize that manipulated ants select certain light levels (~4000 lux), consistent with results of Andriolli et al., 2019, who experimentally found evidence that incident light affects *Ophiocordyceps* fitness. We also found that plots with more open canopies had greater numbers of cadavers producing fruiting bodies. This effect may be directly related to light levels or other aspects of plot habitat linked to canopy openness. Indeed, lighting conditions and circadian rhythms have been hypothesized to affect mechanisms underlying *Ophiocordyceps* manipulation of ants (Das & de Bekker, 2022; de Bekker et al., 2015; de Bekker, Merrow, et al., 2014; de Bekker, Will, et al., 2017; de Bekker & Das, 2022; Trinh et al., 2021; Will et al., 2020). As such, we propose that experiments that manipulate light and other conditions to directly measure their effects on disease outcome would be an appropriate next step.

Additionally, we discovered a non-canonical fungal growth morphology in the brightest graveyard, where the atypical cadavers occupied slightly lower heights than normal cadavers in that plot and across the study. This morphology was characterized by stunted fungal stalks and a dark, dense, thread-like mycelial growth surrounding the cadaver. This growth phenotype may represent a plastic response to increased light levels when adjustment by biting height is not sufficient to place cadavers within a preferred incident light range. The dark mycelium surrounding these cadavers could be a melanized photo-protectant (Cordero & Casadevall, 2017), and the stunted stalks might result from poorer conditions (e.g., increased UV exposure, elevated temperatures, or desiccation). Overall, our findings corroborate previous suggestions that position and timing of cadaver establishment correlate with microclimatic conditions to facilitate proper fungal growth and dispersal (Andersen & Hughes, 2012; Andriolli et al., 2019; Cardoso Neto et al., 2019; Hughes et al., 2011; Lavery et al., 2021).

We also estimated the amount of time needed to produce fruiting bodies in *Ophiocordyceps* manipulated cadavers. Fruiting body formation (25 to 45 days) tended to outpace detachment or loss of cadavers by about 30 days. This pace was somewhat slower than the one to two weeks observed for *O. camponoti-leonardi* in tropical Nakhon Ratchasima, Thailand (Andersen et al., 2009) but much quicker than the 500 to 600 days for *O. kimflemingiae* to overwinter and subsequently fruit in temperate South Carolina, USA (Loreto et al., 2018). *Ophiocordyceps* in Thailand have been suggested to follow an iteroparous reproductive lifestyle, sustaining spore release into the environment over a period of time with high fungal survivorship once fruiting bodies are established (Andersen et al., 2012). At our study sites in Florida, the notable difference in time between fruiting body formation and cadaver detachment may reflect a similar lifestyle, but overall, latitudinal gradients in reproductive strategies may exist within the genus *Ophiocordyceps*.

Fungal mycoparasites can colonize *Ophiocordyceps* tissue emerging from cadavers and likely reduce *Ophiocordyceps* success by inhibiting fruiting body formation or function, depleting resources, or killing *Ophiocordyceps* (Andersen et al., 2012; Araújo et al., 2022 in press; Evans, 1982). Our results support this hyperparasitic interaction for *O. camponoti-floridani*, which had greatly reduced fruiting body formation rates when infected by mycoparasites. Moreover, *N. coronatus* often entirely covered *O. camponoti-floridani* fruiting bodies (Araújo et al., 2022 in press), which may severely limit *O. camponoti-floridani* transmission by creating a physical barrier to spore dispersal.

*Camponotus floridanus* has a caste-based social system in which individuals have plastic behavioral repertoires that reflect molecular, cellular, and social drivers (Das & de Bekker, 2022; Simola et al., 2016; Tripet & Nonacs, 2004). Most infected ants found in our study were minor workers, presumably of the exploratory forager caste. While rare, we did find infected individuals of the morphologically distinct major caste. Differences in infection rates between castes may reflect several factors, including relative caste abundance and exposure to infectious spores, increased immune investment in major ants, and the interaction of caste-specific behavioral profiles with fungal manipulation mechanisms. We see this pattern as an interesting direction for future research.

Many research avenues have yet to be fully explored in understanding the environmental and temporal dynamics of *Ophiocordyceps*-manipulated ant behavior and cadaver development. This study adds to the growing evidence underpinning hypotheses that environmental variables, such as light, weather, and vegetation, affect where cadavers appear. These effects speak to the ways that this fascinating host-pathogen interaction is maintained and may differ across environmental contexts. Although our study included hundreds of ant cadavers monitored over approximately a year, the patterns we observed could be further tested for generalizability at other study sites, ideally for even longer periods of time. Multi-year studies, while demanding, offer some of the best data to disentangle acute environmental responses and seasonal timing – which our current study cannot fully resolve. Further empirical testing is also needed to better understand if and how manipulated ants actively identify and pursue summiting positions at an optimal location. Although we found effects for vegetation type and light levels in relation to relative cadaver abundance, more detailed continuous observations and experimentation are required to understand if these traits are adaptive to *Ophiocordyceps* fitness, and how this varies across *Ophiocordyceps* species and geographic locations. Similarly, hypothesis driven experiments or detailed surveys of microhabitat use are approachable next steps to dissect the correlation of weather patterns and cadaver accumulation. We envision that both controlled laboratory tests of specific environmental factors and experimental translocations of cadavers under natural field conditions would be fruitful endeavors to undertake. Studies with Floridian “zombie ants” may be especially amenable to field translocations, as most cadavers are found on epiphytic *Tillandsia*, allowing simple exchanges of cadaver positions without killing the bitten plant. Furthermore, with this study we add our voices to calls for more “boots on the ground” natural history and detailed, high-replication field observations (Wilson, 2017). With the advent of modern (molecular) biology, field studies became somewhat underappreciated. However, such reports remain foundational efforts that offer both insights into the drivers of variation and adaptations across natural systems and hypotheses based on real-world data. Of course, hand in hand with such efforts, finely focused laboratory experiments would be necessary if one aims to answer mechanistic questions. Therefore, the integration of laboratory and field experiments with molecular biology would have the power to reveal the drivers of interactions and selective processes that affect the co-evolved symbioses between *Ophiocordyceps* fungi, their mycoparasites, and their ant hosts.

## ACKNOWLEDGEMENTS

We thank the Florida Department of Agriculture and Consumer Service’s Florida Forest Service and the Seminole County’s Leisure Services Department, Greenways and Natural Lands Division for permitting our field observations and collections. We thank Davide Dal Pos for helping with the macrophotography of representative samples, Brittany Lebert for obtaining sequence data from one of the field samples, and Biplabendu Das, Brianna Santamaria, Meghan Flynn, and Hanniah Myers for assisting in the field. Two anonymous reviewers offered constructive comments to improve the manuscript, for which we are appreciative. The work presented in this manuscript has been supported by startup funds from the University of Central Florida, made available to Charissa de Bekker. Charissa de Bekker is also supported by NSF CAREER 1941546.

## DATA AVAILABILITY

DNA sequence data for *O. camponoti-floridani* have been deposited under GenBank ON495684. Field and weather station data have been deposited on Dryad (https://doi.org/10.5061/dryad.4tmpg4fd8).

## SUPPORTING INFORMATION

Additional supporting information may be found in the online version of this article.

Figure S1 Study sites and vegetation.

Figure S2 Relative canopy openness does not clearly correlate with individual cadaver incident light.

Figure S3 Visit intervals spanning observation of a cadaver fruiting body or detachment.

Figure S4 Visit intervals spanning observation of a cadaver takeover by mycoparasites and monthly occurrence.

Table S1 Cadaver accumulation by vegetation model overview and ranking by AIC.

Table S2 Each effect per cadaver accumulation by vegetation model more efficient than the null is given with a summary of statistics.

Table S3 Cadaver height by light model overview and ranking by AIC.

Table S4 Each effect per cadaver height by light model more efficient than the null model is given with a summary of statistics.

Table S5 Cadaver accumulation rate by weather and habitat model overview and ranking by AIC.

Table S6 Each effect in the top five cadaver accumulation rate by weather and habitat models is given with a summary of statistics.

Table S7 Cadaver transmission outcome by weather and habitat model overview and ranking by AIC.

Table S8 Each effect for the top five cadaver accumulation by vegetation models is given with a summary of statistics.

**Supplemental Figure 1:**
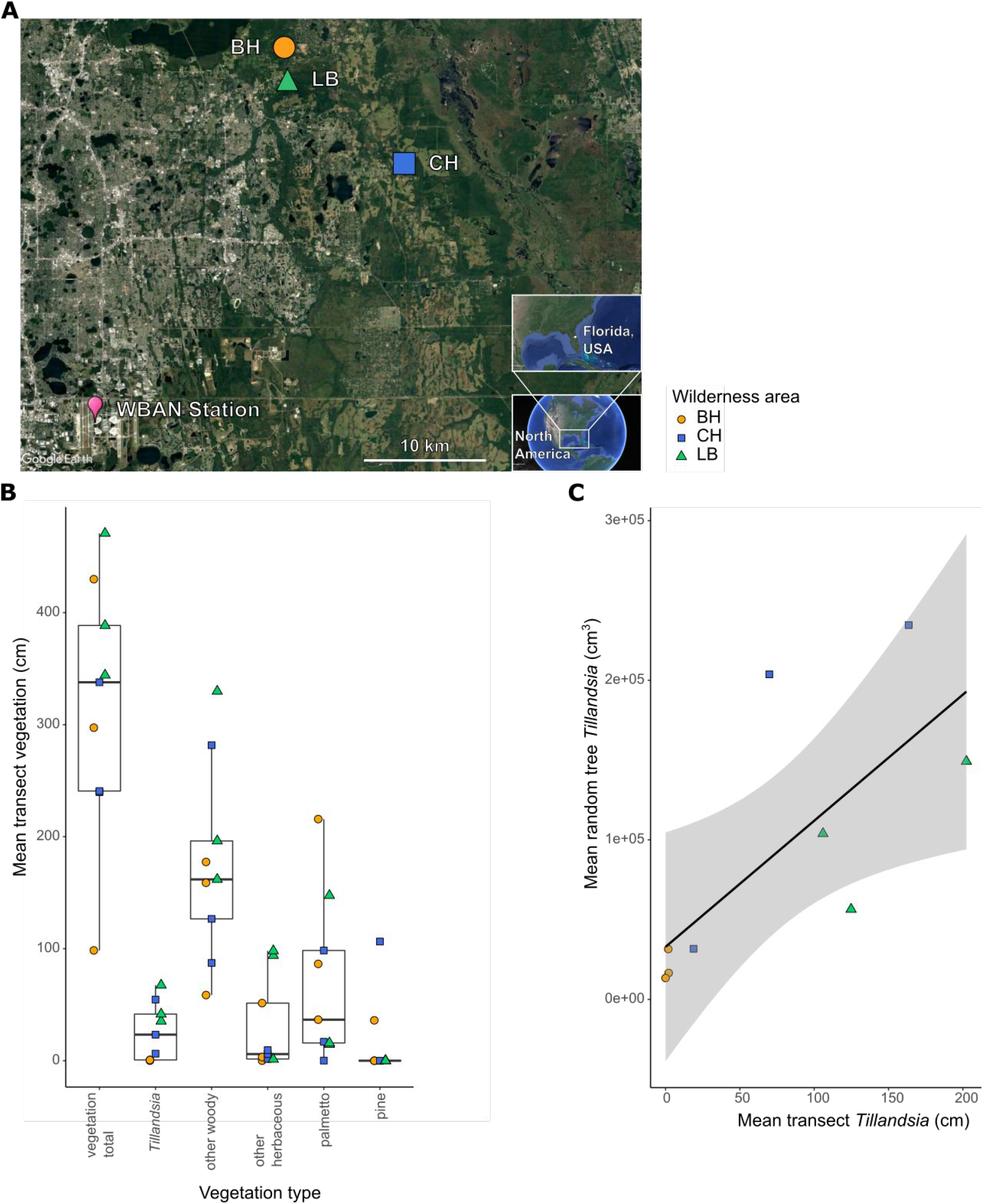
Study sites and vegetation. A) We established a total of nine graveyard plots across three wilderness areas in Seminole Co., Orlando, FL, USA. The graveyards within a given wilderness area were on average 0.39 km apart from one another. The NOAA weather station WBAN: 12815 (magenta drop point) provided climate data for all graveyards and was ca. 32 km from our study sites. Image produced with Google Earth Pro (v7.3.4.8248). B) Transect vegetation measurements at each graveyard show that our field sites captured the variation of Central Florida mesic hammock habitat in which *Ophiocordyceps-infected* ants are generally found. Only a few graveyards appear to be possible outliers for certain types of vegetation. C) A linear model (black line) (R^2^ = 0.53) describing the volume of *Tillandsia* counted on trees within a graveyard by transect measurements of *Tillandsia* in that graveyard, suggesting that both methods are in general agreement with regards to estimating the coverage of *Tillandsia* abundance. The shaded area represents a 95% confidence interval. In all figures Black Hammock Wilderness (BH) graveyards are indicated as orange circles, Chuluota Wilderness (CH) as blue squares, and Little Big Econ State Forest (LB) as green triangles.

**Supplemental Figure 2:**
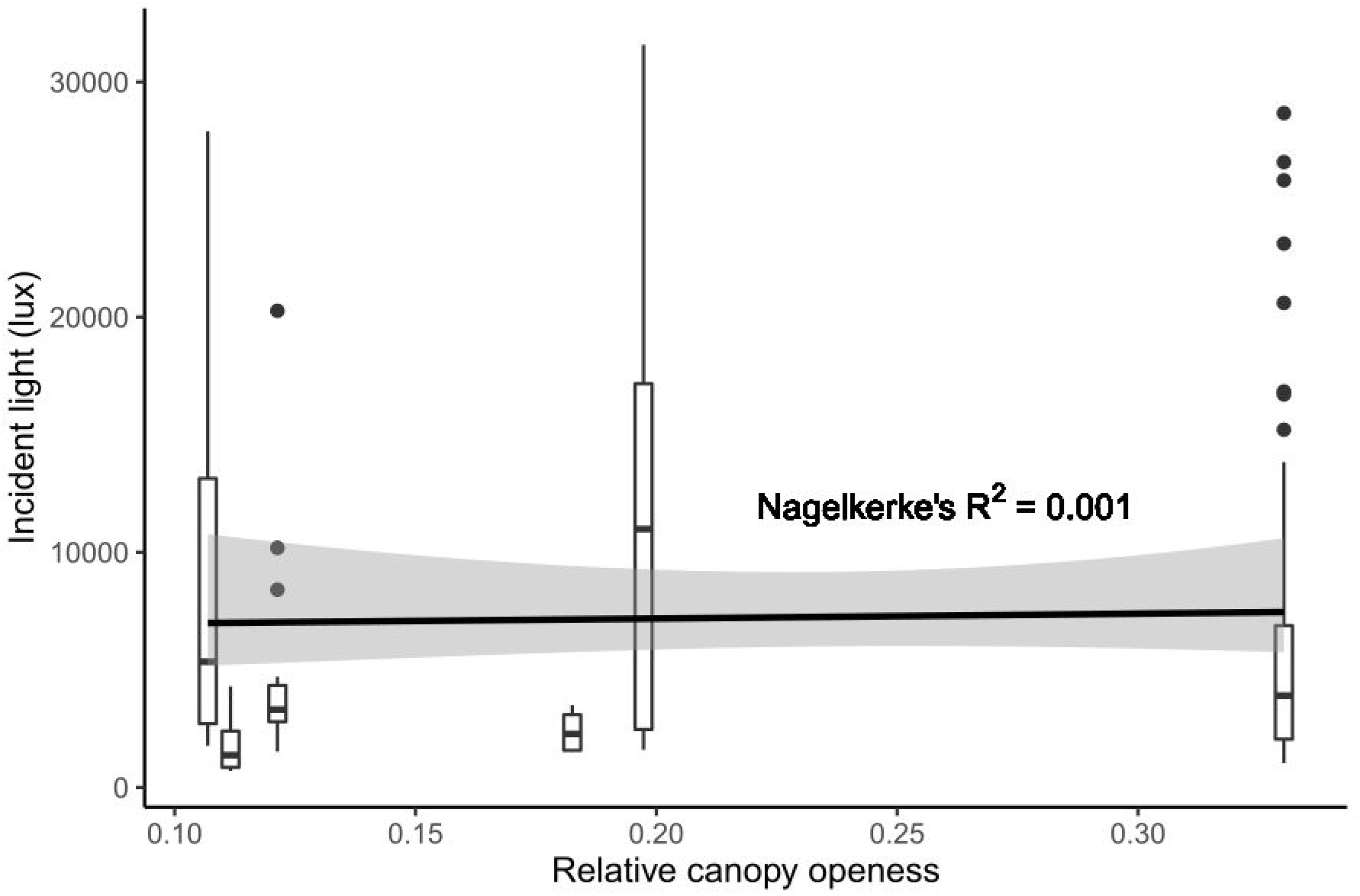
Relative canopy openness does not clearly correlate with individual cadaver incident light. Canopy openness appears to more strongly correlate with cadaver height (Fig 3B) rather than cadaver incident light correlates with height (Fig 3C). Following this, we indeed found that there appears to be a minimal direct relationship between canopy openness and incident light when modeling incident light by canopy openness (Gamma distribution GLM, Nagelkerke’s R^2^ = 0.001, coefficient = −3.8E-6, p = 0.801).

**Supplemental Figure 3:**
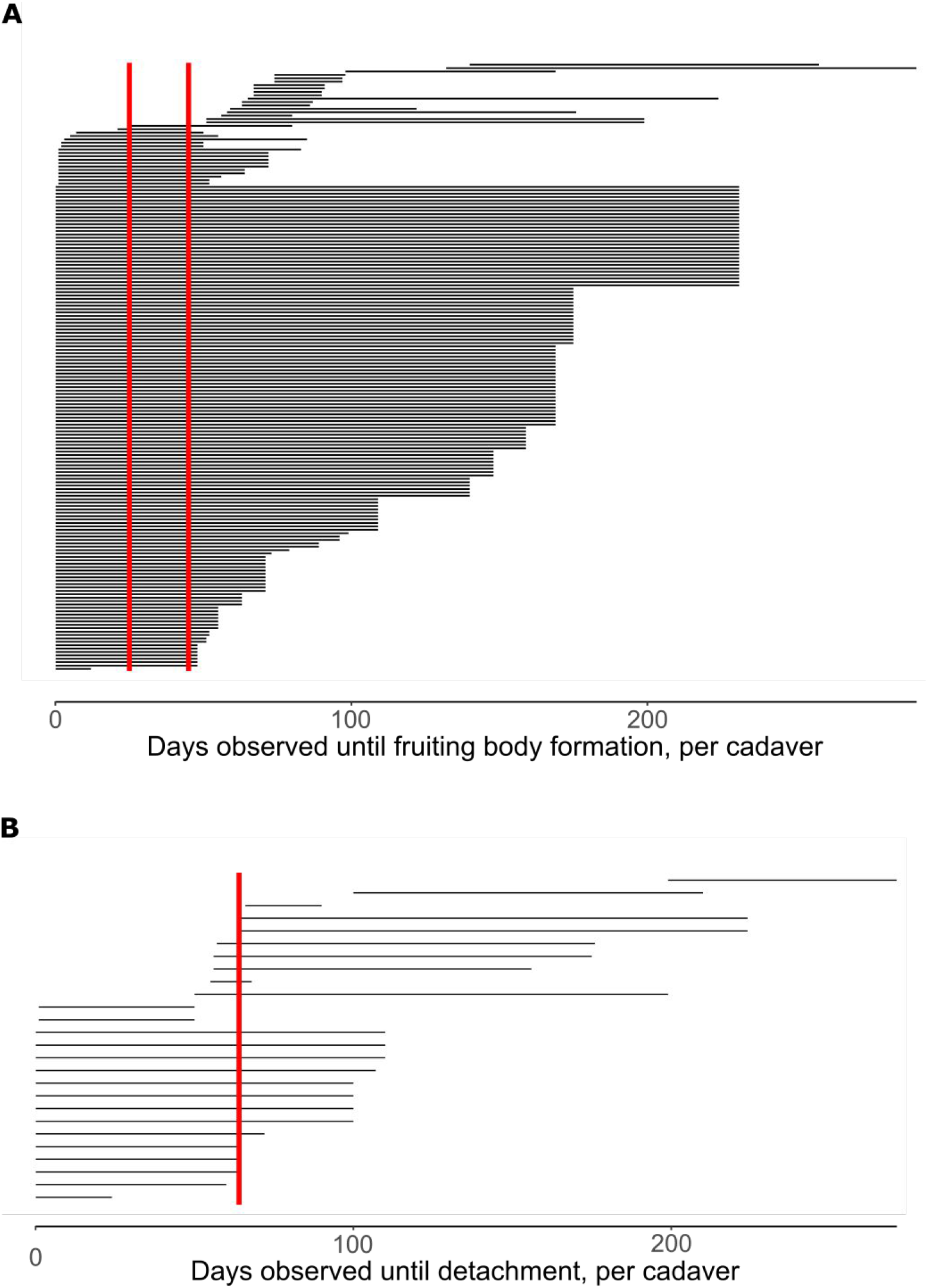
Visit intervals spanning observation of a cadaver fruiting body or detachment. A) Black lines represent a single cadaver, showing the estimated lapsed time between a cadaver’s establishment in a graveyard and the formation of a fruiting body. For cadavers with a minimum time equal to zero days, this means that during the previous visit to that graveyard the cadaver was not found, and then upon its first entry into the study it already had a fruiting body. Red vertical lines indicate a range of days shared by the most observations: 25 – 45 days (n = 160 of 179 fruiting body producing cadavers). B) Black lines represent a single cadaver and show the estimated lapsed time between a cadaver’s establishment in a graveyard and the moment it detached from its perch. For cadavers that detached and had a minimum time equal to zero days, this means that during the previous visit the ant was found for the first time and before the next visit had detached. The red vertical line shows 64 days as the most common time until observation of detachment (n = 19 of 26 detaching cadavers). The typical time needed for fruiting body formation was shorter than the typical time to detachment, suggesting that most established cadavers have an opportunity to transmit.

**Supplemental Figure 4:**
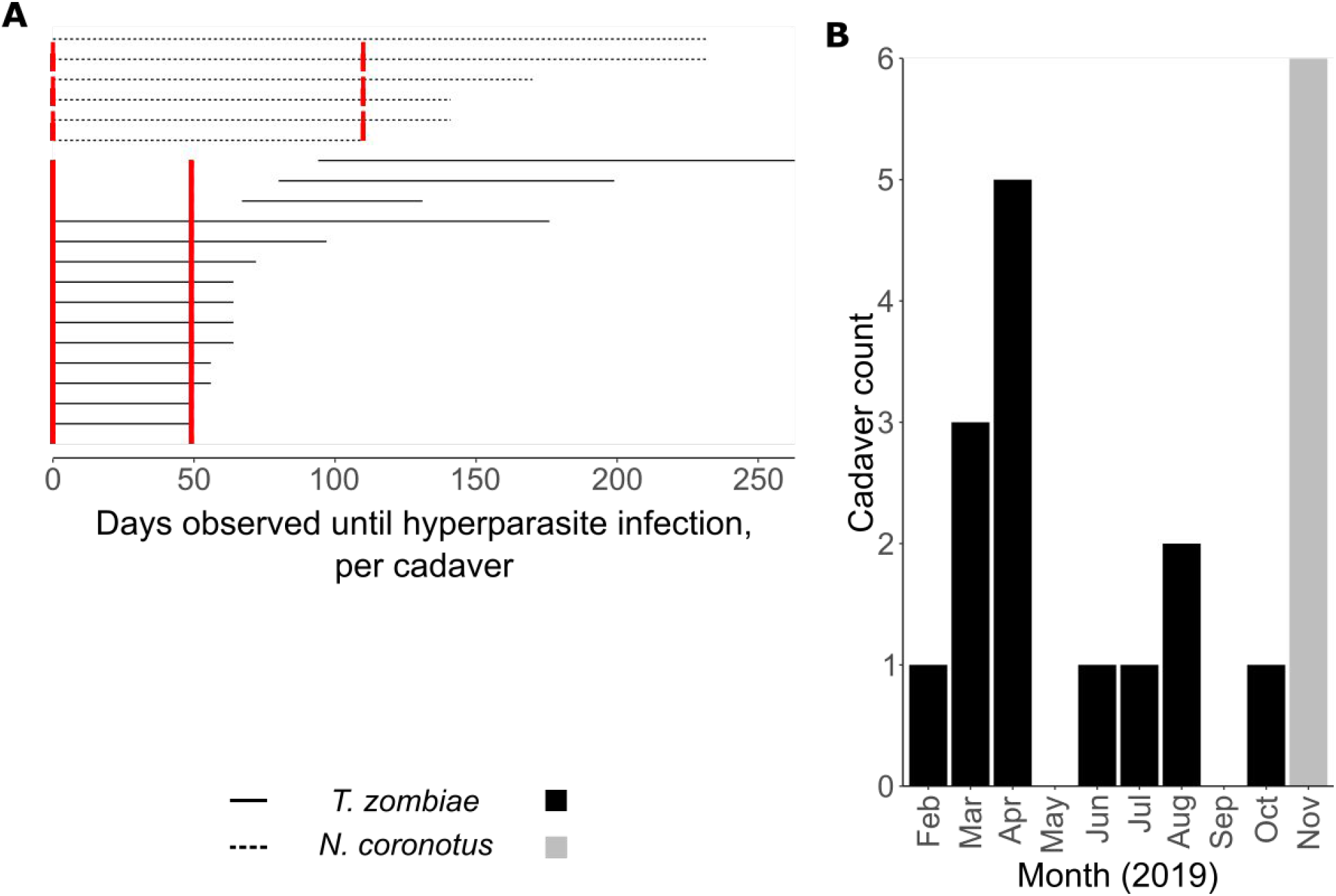
Visit intervals spanning observation of a cadaver takeover by mycoparasites and monthly occurrence. A) Each line represents a single cadaver, showing the estimated lapsed time between a cadaver’s establishment in a graveyard and the infection by a mycoparasite. Solid black lines represent the time until observation of *T. zombiae* infection and dashed black lines represent *N. coronotus* infections. Red lines indicate the highest and lowest most-spanned days until observed mycoparasitism (*T. zombiae* solid lines between 0 – 49 days, *N. coronotus* dashed lines between 0 – 110 days). B) Observations of *T. zombiae* infections were recorded most months of the year, while observation of *N. coronotus* only occurred in November. However, due to the long interval during which *N. coronotus* may have colonized, we cannot make a statement about possible seasonality.

**Supplemental Table 1:** Cadaver accumulation by vegetation model overview and ranking by AIC. We selected model “Full” as the final model. Variables shown: vegetation type (veg_type), relative abundance of vegetation (veg%), graveyard plot (plot), and wilderness area (area), with additive effects (+), interactive effects (*), and nested effects (/).

| Model name | Fixed effects | Random effects | AIC | R<sup>2</sup> conditional | R<sup>2</sup> marginal |
| --- | --- | --- | --- | --- | --- |
| Full | veg_type*veg% | area/plot | 289.33 | 0.640 | 0.548 |
| Additive | veg_type + veg% | park | 395.40 | 0.525 | 0.525 |
| Type only | veg_type + plot | NA | 442.82 | NA | 0.511 |
| null | 1 | NA | 1283.05 | NA | 0.00 |

**Supplemental Table 2:** Each effect per cadaver accumulation by vegetation model more efficient than the null is given with a summary of statistics. Variables shown: vegetation type (veg_type), relative abundance of vegetation (veg%), graveyard plot (plot), wilderness area (area), and interaction between variables (:). Categorical fixed effects are shown for each level except the intercept level, “other woody” for veg_type and LB1 for plot). Standard error is SE and standard deviation is SD.

| Effect | Coefficient (fixed) or<br>variance (random) | SE (fixed) or<br>SD (random) | z value | p value |
| --- | --- | --- | --- | --- |
| **Full model** |  |  |  |  |
| veg_type – *Tillandsia* | 1.43 | 0.59 | 2.42 | 0.016 |
| veg_type – herbaceous | -1.53 | 0.71 | -2.15 | 0.032 |
| veg_type – palmetto | -2.13 | 1.28 | -1.66 | 0.096 |
| veg_type – pine | -0.57 | 0.64 | -0.89 | 0.373 |
| veg% | 1.38 | 1.07 | 1.29 | 0.197 |
| *Tillandsia*:veg% | 30.45 | 3.09 | 9.86 | < 0.001 |
| herbaceous:veg% | 4.85 | 3.15 | 1.54 | 0.122 |
| palmetto:veg% | -1.17 | 3.15 | -0.37 | 0.710 |
| pine:veg% | 5.47 | 1.47 | 3.72 | <0.001 |
| plot:area (random) | 0.38 | 0.62 | NA | NA |
| park (random) | 0.45 | .067 | NA | NA |
| **Additive model** |  |  |  |  |
| veg_type – *Tillandsia* | 4.15 | 0.35 | 11.90 | < 0.001 |
| veg_type – herbaceous | -0.27 | 0.43 | -0.63 | 0.530 |
| veg_type – palmetto | -1.77 | 0.50 | -3.58 | < 0.001 |
| veg_type – pine | 0.89 | 0.33 | 2.67 | 0.008 |
| veg% | 2.87 | 0.56 | 5.11 | < 0.001 |
| park (random) | < 0.001 | 0.02 | NA | NA |
| **Type only model** |  |  |  |  |
| veg_type – *Tillandsia* | 2.70 | 0.17 | 15.79 | < 0.001 |
| veg_type – herbaceous | -1.72 | 0.31 | -5.51 | < 0.001 |
| veg_type – palmetto | -2.69 | 0.47 | -5.74 | < 0.001 |
| veg_type – pine | -0.40 | 0.21 | -1.93 | 0.053 |
| plot – LB2 | < 0.001 | 0.33 | < 0.001 | 1.000 |
| plot – LB3 | < 0.001 | 0.26 | < 0.001 | 1.000 |
| plot – BH1 | < 0.001 | 0.26 | < 0.001 | 1.000 |
| plot – BH2 | < 0.001 | 0.31 | < 0.001 | 1.000 |
| plot – BH3 | < 0.001 | 0.37 | < 0.001 | 1.000 |
| plot – CH1 | < 0.001 | 0.28 | < 0.001 | 1.000 |
| plot – CH2 | < 0.001 | 0.25 | < 0.001 | 1.000 |
| plot – CH3 | < 0.001 | 0.32 | < 0.001 | 1.000 |

**Supplemental Table 3:** Cadaver height by light model overview and ranking by AIC. We selected model “Canopy” as the final model. Variables shown: percent canopy openness (canopy), incident light (lux), graveyard plot (plot), and wilderness area (area), with additive effects (+).

| Model name | Fixed effects | Random effects | AIC | Nagelkerke’s R<sup>2</sup> |
| --- | --- | --- | --- | --- |
| Canopy | canopy + area | NA | 1161.99 | 0.144 |
| Plot | plot | NA | 1163.60 | 0.165 |
| Canopy & incident | canopy + lux + area | NA | 1163.84 | 0.146 |
| null | 1 | NA | 1171.36 | < 0.001 |

**Supplemental Table 4:** Each effect per cadaver height by light model more efficient than the null model is given with a summary of statistics. Variables shown: percent canopy openness (canopy), incident light (lux), graveyard plot (plot), and wilderness area (area). Categorical fixed effects are shown for each level except the intercept level, LB1 for plot and BH for area. Standard error is SE and standard deviation is SD.

| Effect | Coefficient (fixed) or<br>variance (random) | SE (fixed) or<br>SD (random) | t value | p value |
| --- | --- | --- | --- | --- |
| **Canopy** |  |  |  |  |
| canopy | -0.37 | 0.15 | -2.40 | 0.018 |
| area – CH | 0.36 | 0.30 | 1.19 | 0.236 |
| area – LB | -0.18 | 0.16 | -1.11 | 0.269 |
| **Plot** |  |  |  |  |
| plot – LB3 | -0.02 | 0.29 | -0.05 | 0.96 |
| plot – BH1 | -0.10 | 0.20 | -0.51 | 0.61 |
| plot – BH2 | 0.32 | 0.21 | 1.59 | 0.12 |
| plot – BH3 | 0.09 | 0.26 | 0.36 | 0.72 |
| plot – CH2 | -0.22 | 0.19 | -1.18 | 0.24 |
| **Canopy & incident** |  |  |  |  |
| canopy | -0.35 | 0.16 | -2.21 | 0.029 |
| lux | -0.02 | 0.05 | -0.42 | 0.679 |
| area – CH | 0.33 | 0.31 | 1.04 | 0.300 |
| area – LB | -0.18 | 0.16 | -1.13 | 0.263 |

**Supplemental Table 5:** Cadaver accumulation rate by weather and habitat model overview and ranking by AIC. We selected model “sigW-canopy” as the final model. Variables shown: mean daily maximum temperature (T), mean daily precipitation (P), mean daily relative humidity (H), percent canopy openness (canopy), random-tree *Tillandsia* abundance (Till), line-intersect *Tillandsia* relative abundance (Till%), and graveyard plot (plot), with additive effects (+), interactive effects (*), and interaction-only effects (:). sigW indicates a set of weather terms that were significant from a full model containing a three-way interaction and addW indicates an additive-only set of all weather effects.

| Model name | Fixed effects | Random effects | AIC | R<sup>2</sup> marginal | R<sup>2</sup> conditional |
| --- | --- | --- | --- | --- | --- |
| sigW-canopy | T:P:H + T + P + openness% | plot | 271.77 | 0.56 | 0.75 |
| sigW-Till | T:P:H + T + P + Till | plot | 272.70 | 0.59 | 0.78 |
| sigW-Till% | T:P:H + T + P + Till% | plot | 274.00 | 0.57 | 0.78 |
| sigW | T:P:H + T + P | plot | 275.17 | 0.40 | 0.76 |
| W-Till%-canopy | T*P*H + Till% + openness% | plot | 275.72 | 0.68 | 0.79 |
| W-canopy | T*P*H + openness% | plot | 276.27 | 0.58 | 0.76 |
| W-Till | T*P*H + Till | plot | 276.57 | 0.64 | 0.79 |
| W-Till% | T*P*H + Till% | plot | 277.86 | 0.62 | 0.80 |
| W-Till-canopy | T*P*H + Till + openness% | plot | 276.70 | 0.65 | 0.78 |
| W | T*P*H | plot | 279.69 | 0.43 | 0.78 |
| addW-canopy | T + P + H + openness% | plot | 279.86 | 0.42 | 0.70 |
| addW-Till%-canopy | T + P + H + Till% + openness% | plot | 280.91 | 0.46 | 0.71 |
| addW-Till | T + P + H + Till | plot | 281.11 | 0.40 | 0.72 |
| addW-Till-canopy | T + P + H + Till + openness% | plot | 281.47 | 0.43 | 0.70 |
| addW-Till% | T + P + H + Till% | plot | 281.72 | 0.39 | 0.73 |
| null | 1 | plot | 299.93 | NA | 0.62 |

**Supplemental Table 6:** Each effect in the top five cadaver accumulation rate by weather and habitat models is given with a summary of statistics. Variables shown: mean daily maximum temperature (T), mean daily precipitation (P), mean daily relative humidity (H), percent canopy openness (canopy), random-tree *Tillandsia* abundance (Till), line-intersect *Tillandsia* relative abundance (Till%), and graveyard plot (plot), with additive effects (+), and interactions between variables (:). Standard error is SE and standard deviation is SD.

| Effect | Coefficient (fixed) or<br>variance (random) | SE (fixed) or<br>SD (random) | z value | p value |
| --- | --- | --- | --- | --- |
| **sigW-canopy** |  |  |  |  |
| T | 0.28 | 0.07 | 3.95 | < 0.001 |
| P | -0.27 | 0.09 | -2.90 | 0.004 |
| T:P:H | 0.20 | 0.07 | 2.99 | 0.003 |
| canopy | 0.26 | 0.10 | 2.71 | 0.007 |
| plot (random) | 0.07 | 0.27 | NA | NA |
| **sigW-Till** |  |  |  |  |
| T | 0.27 | 0.07 | 3.84 | < 0.001 |
| P | -0.26 | 0.09 | -2.86 | 0.004 |
| T:P:H | 0.21 | 0.07 | 3.03 | 0.002 |
| Till | 0.24 | 0.10 | 2.44 | 0.015 |
| plot (random) | 0.08 | 0.29 | NA | NA |
| **sigW-Till%** |  |  |  |  |
| T | 0.27 | 0.07 | 3.82 | < 0.001 |
| P | -0.26 | 0.09 | -2.79 | 0.005 |
| T:P:H | 0.20 | 0.07 | 2.90 | 0.004 |
| Till% | 0.23 | 0.12 | 1.95 | 0.051 |
| plot (random) | 0.10 | 0.32 | NA | NA |
| **sigW** |  |  |  |  |
| T | 0.27 | 0.07 | 3.85 | < 0.001 |
| P | -0.25 | 0.09 | -2.72 | 0.007 |
| T:P:H | 0.18 | 0.07 | 2.70 | 0.007 |
| plot (random) | 0.15 | 0.39 | NA | NA |
| **W-Till%-canopy** |  |  |  |  |
| T | 0.36 | 0.09 | 2.89 | < 0.001 |
| P | -0.24 | 0.11 | -2.22 | 0.026 |
| H | -0.06 | 0.14 | -0.42 | 0.672 |
| T:P | -0.25 | 0.14 | -1.85 | 0.064 |
| T:H | 0.06 | 0.16 | 0.39 | 0.696 |
| P:H | -0.10 | 0.15 | -0.65 | 0.513 |
| T:P:H | 0.38 | 0.13 | 2.88 | 0.004 |
| Till% | 0.17 | 0.10 | 1.73 | 0.084 |
| canopy | 0.21 | 0.09 | 2.29 | 0.022 |
| plot (random) | 0.05 | 0.23 | NA | NA |

**Supplemental Table 7:**
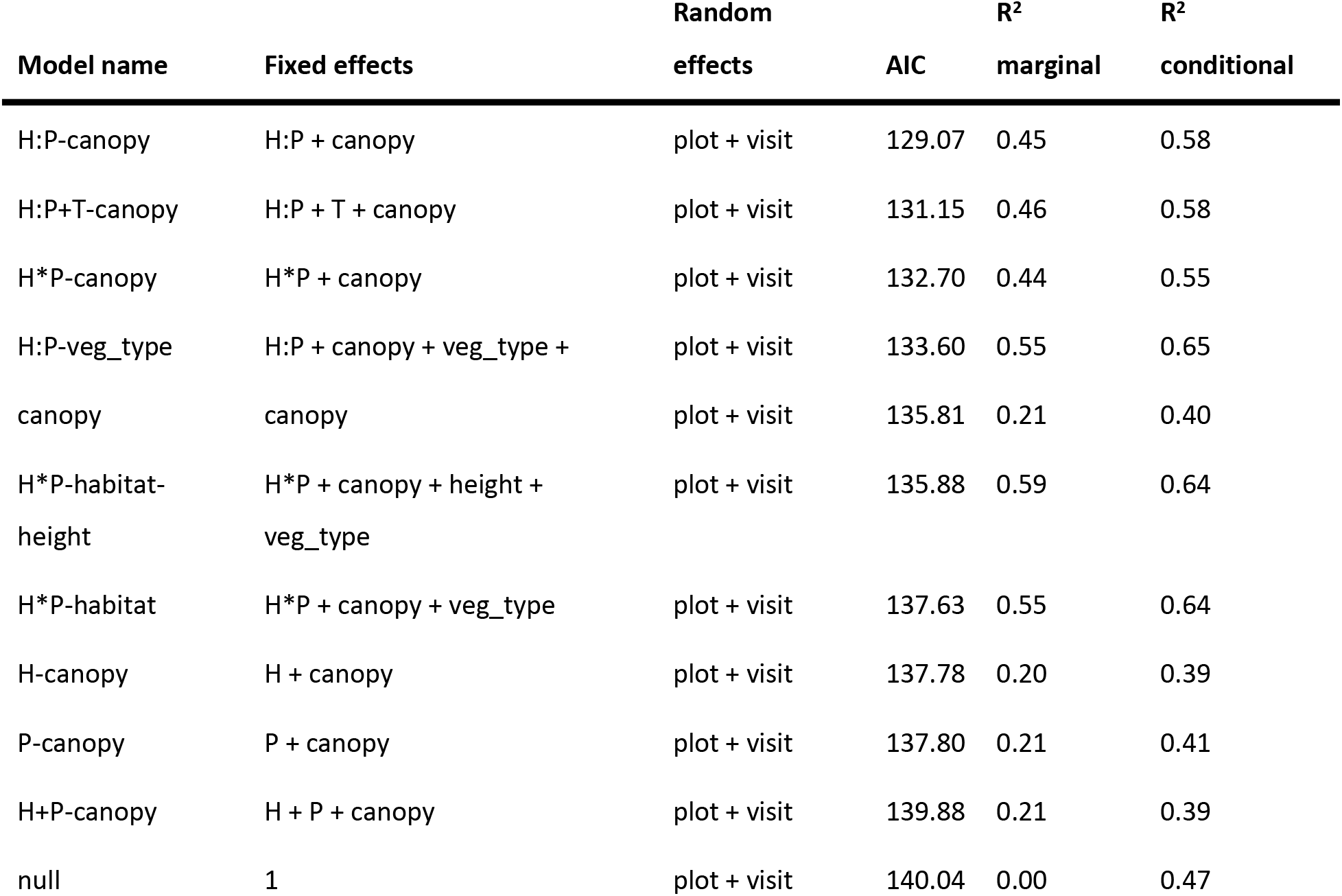
Cadaver transmission outcome by weather and habitat model overview and ranking by AIC. We selected model “H:P-canopy” as the final model. Variables shown: mean daily relative humidity (H), mean daily precipitation (P), vegetation type (veg_type), percent canopy openness (canopy), cadaver height (height), graveyard plot (plot), and visit time bin (visit), with additive effects (+), interactive effects (*), and interaction-only effects (:).

**Supplemental Table 8:**
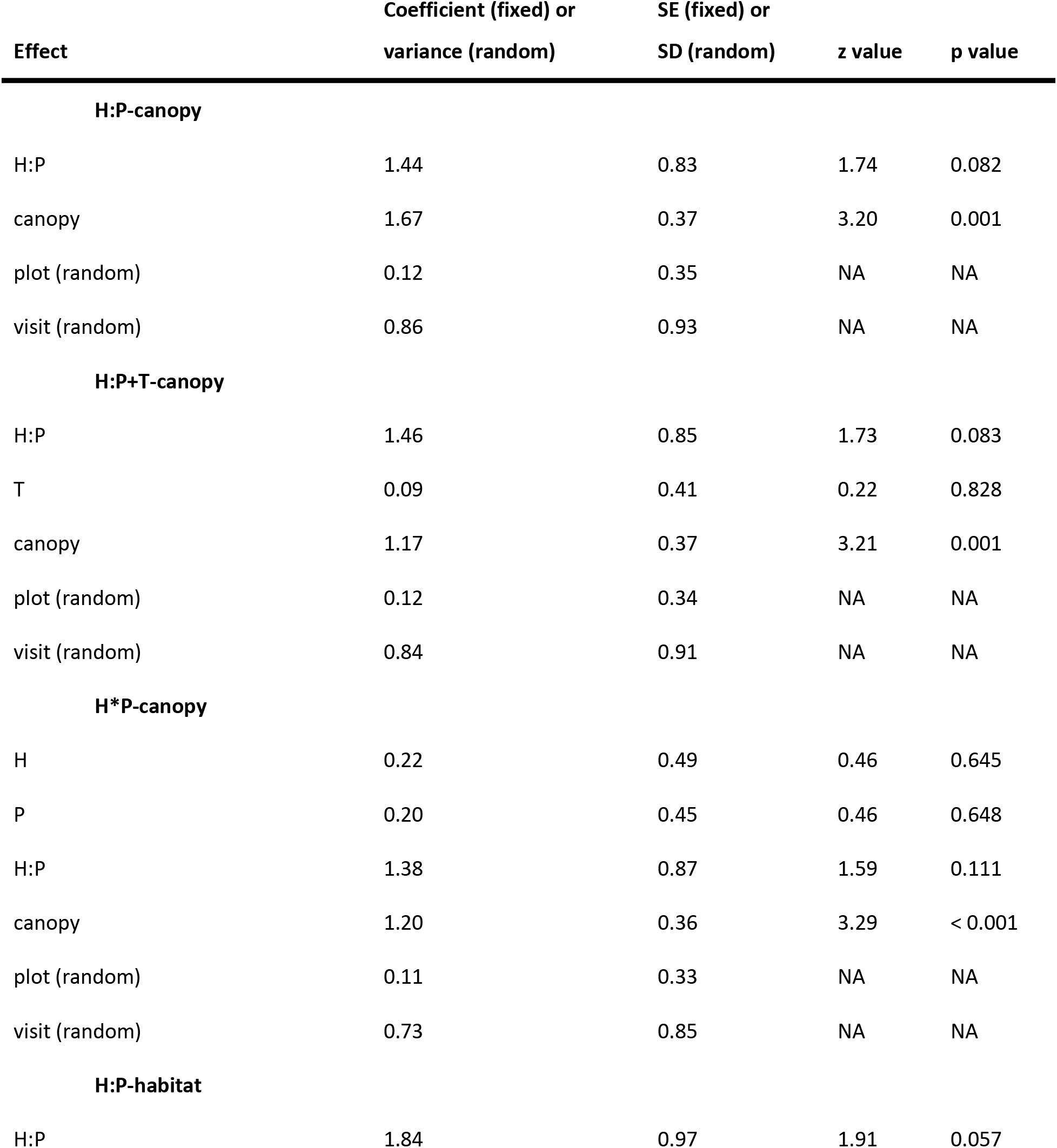
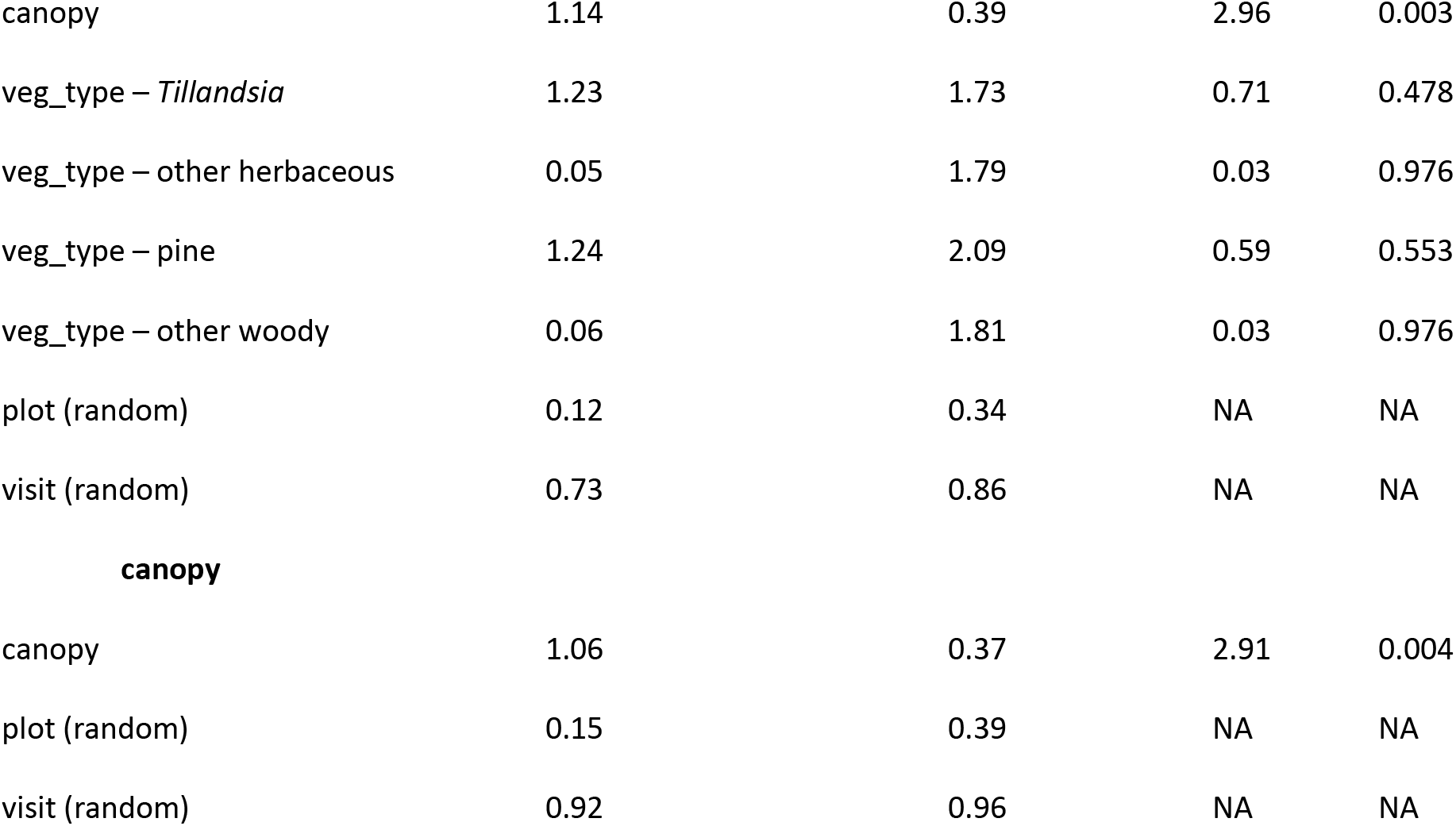
Each effect for the top five cadaver accumulation by vegetation models is given with a summary of statistics. Variables shown: mean daily relative humidity (H), mean daily precipitation (P), vegetation type (veg_type), percent canopy openness (canopy), graveyard plot (plot), and visit time bin (visit), with, and interaction of effects (:). Categorical fixed effects are shown for each level except the intercept level (palmetto for veg_type). Standard error is SE and standard deviation is SD.

